# Making Strides in Doctoral-Level Career Outcomes Reporting: Surveying the Landscape of Classification and Visualization Methodologies and Creating a Crosswalk Tool

**DOI:** 10.1101/2021.07.12.451657

**Authors:** Tammy Collins, Deepti Ramadoss, Rebekah L. Layton, Jennifer MacDonald, Ryan Wheeler, Adriana Bankston, Abigail Stayart, Yi Hao, Jacqueline N Robinson-Hamm, Melanie Sinche, Scott Burghart, Aleshia Carlsen-Bryan, Pallavi Eswara, Heather Krasna, Hong Xu, Mackenzie Sullivan

## Abstract

The recent movement underscoring the importance of career taxonomies has helped usher in a new era of transparency in PhD career outcomes. The convergence of discipline-specific organizational movements, interdisciplinary collaborations, and federal initiatives have all helped to increase PhD career outcomes tracking and reporting. Transparent and publicly available PhD career outcomes are being used by institutions to attract top applicants, as prospective graduate students are factoring these in when deciding on the program and institution in which to enroll for their PhD studies. Given the increasing trend to track PhD career outcomes, the number of institutional efforts and supporting offices for these studies have increased, as has the variety of methods being used to classify and report/visualize outcomes. This report identifies and summarizes currently available PhD career taxonomy tools, resources, and visualization options to help catalyze and empower institutions to develop and publish their own PhD career outcomes. Similar fields between taxonomies were mapped to create a new crosswalk tool. This work serves as an empirical review of the career outcome tracking systems available and highlights organizations, consortia, and funding agencies that are impacting policy change toward greater transparency in PhD career outcomes reporting.

## INTRODUCTION

In the past decade, there have been growing calls to action for institutions to collect and disseminate career outcomes data for graduate students and postdocs, and to develop common standards for reporting these data^1–7^ including the National Institutes of Health Biomedical Research Workforce Working Group Report 2012^8^. These calls are linked to broad systemic issues that are well-documented^9^, including a highly competitive faculty job market with far fewer available positions relative to the supply of PhDs, compensation and training length concerns for postdoctoral scholars, and changing educational and career interests of PhDs.

Numerous efforts and approaches to address the need for better career outcomes data collection have emerged, many of which are described in this report. Efforts coalesce into three major approaches: building coalitions, updating funding obligations, and promoting transparent career outcomes. First, the formation of coalitions of stakeholder working groups or institutions committed to common standards has created purpose-driven communities of thought and action. These groups have clarified the central issues and concentrated the call to action, exemplified by the creation and adoption of the Unified Career Outcomes Taxonomy (UCOT)^7^. Some of these groups include Rescuing Biomedical Research (http://rescuingbiomedicalresearch.org/), the NIH Broadening Experiences in Scientific Training Consortium (https://commonfund.nih.gov/workforce), the Coalition for Next Generation Life Science (https://nglscoalition.org/), and topically-focused meetings such as the Future Of Bioscience Graduate and Postdoctoral Training conference (FOBGAPT 1 & FOBGAPT 2; https://gs.ucdenver.edu/fobgapt2/main.php). A second set of efforts have focused on updating prerequisites to funding to require the collection and dissemination of institutional outcomes data (such as the National Institute of General Medical Sciences’ Request for Applications (NIGMS RFA) requirements for T32 Training Grants). An increasingly common third approach has focused on the development and implementation of institute- or discipline-specific practices for publicly sharing outcomes data, exemplified by recent activities from the American Historical Association (AHA; https://www.historians.org/wherehistorianswork) and the National Institute of Environmental Health Sciences^10^ (NIEHS; https://www.niehs.nih.gov/careers/research/fellows/alumni-outcomes/index.cfm), thereby changing the standard expectation for other professional societies and insitutions. These efforts to collect, assess, and publish career outcomes of PhD graduates are becoming standard practice and carry significant benefit to institutions. Internally, the data can be used to inform curricular, training, budgetary, benchmarking, and recruitment priorities, while current and prospective trainees might use the data to make informed strategic decisions about their career choices and preparation. On the scale of the global workforce, transparent and standardized reporting of career outcomes data clarifies the PhDs’ prevalence and impact on society.

Our aim is to decrease the barriers for institutions to collect and report on the career outcomes for their graduate students by summarizing the various options and resources for undertaking these important tasks, and highlighting their key features so that informed decisions can be made about which tools best suit a particular institution’s needs. This report describes: 1) institutions or groups with clearly defined or widely used taxonomies or classification systems; 2) methods for visualizing PhD career outcomes; and 3) tools, resources, or organizations that assist constituents in collecting and reporting this information. We also describe the development of a crosswalk tool^11^, in which similar fields among all the taxonomies examined were mapped to showcase their commonalities.

## METHODS

We conducted a review of graduate career outcomes taxonomies that have been either used, developed, or published within the past five years. Preference for inclusion was given to taxonomies that met one of the following criteria: 1) widely used at a national level; 2) developed by consensus with multiple stakeholders, including those at professional societies; or 3) those with clearly defined categories and rubrics that facilitate reproducibility. This resulted in the identification of 13 taxonomies, which were developed by governmental organizations, professional societies, Universities and Consortia of institutions, etc. Preference was also given for taxonomies that contained classification of doctoral-level career outcomes (either developed specifically for them or used for doctoral populations), while some may also have applicability to those with Master’s degrees. Descriptions and examples of each taxonomy in action were collected and described herein.

Visualizations were selected in part by reviewing the Graduate Career Consortium Outcomes Database^12^ and showcasing examples that depict multi-dimensional outcomes in innovative, meaningful ways. Resources, tools, and coalitions were selected for inclusion by reviewing published reports (including conference abstracts), literature, and websites from within the past five years— with information more commonly being found from within the past three. All of the referenced websites contained within this manuscript link to beneficial resources, and it must be noted that website addresses are prone to change. The links shown herein are current as of June, 2021.

To create the taxonomy crosswalk table, we mapped relative equivalencies between the taxonomies described herein as well as the classification methodologies of 17 additional groups, including public and private universities, consortia, or professional societies. In mapping these taxonomies, a number of challenges occurred. For example, some taxonomies were too comprehensive to fully map within the table developed (e.g., there were nearly 1500 categories to choose from), and these omissions were noted within the table. Some categories had a tally higher than the total number of taxonomies examined because they were present in multiple ways within a single taxonomy (e.g., tenure-track faculty may have appeared as a variety of different professor job titles). Additionally, some categories were repeated for the purposes of alignment; an asterisk (*) was used to indicate when this “one-to-many” mapping occurred. Another key challenge is that no two taxonomies have categories that are 100% equivalent. This was especially apparent when examining employment categorization between different countries. Nevertheless, efforts were made to ascertain the fundamental meaning of each data field in order to best highlight approximate equivalencies between taxonomies. Furthermore, in order to prevent the loss of granularity when aligning taxonomies that are more complex, multiple rows are depicted back-to-back with the same color to highlight categories that are related.

### 1. SYSTEMS FOR CLASSIFYING CAREER OUTCOMES

Below, we highlight a variety of taxonomies/classification systems that have been used within the past decade. The systems are loosely ordered and grouped based upon whether one builds on another, and whether: 1) it is a nationally-based survey; 2) it was developed by an individual institution or consortium; or 3) it was created by a professional association.

#### NATIONAL SURVEY-BASED SYSTEMS

##### National Science Foundation (NSF) Survey of Earned Doctorates (SED)

###### Basic Definition

Since 1957, the National Science Foundation (NSF) has administered an annual census-type survey to all research doctorate earners from accredited U.S. institutions, titled the Survey of Earned Doctorates (SED) (https://nsf.gov/statistics/srvydoctorates/#qs). Data are reported at the end of each calendar year following the survey administration date. This survey is administered using a cross-sectional design to capture information about graduate training and education, and includes information about career outcomes. Data are collected directly from PhD graduates. Development of this tool was sponsored by the National Center for Science and Engineering Statistics (NCSES) (https://www.nsf.gov/statistics/) within the NSF, along with multiple federal organizations, including the National Institutes of Health, Department of Education, and the National Endowment for the Humanities, to provide national-level data and reports on outcomes of doctoral training. The SED provides an annual snapshot of the first destinations of doctoral degree recipients.

###### Development

The SED contains information about educational and training history, and asks graduates to choose from a set of options regarding what best describes their post PhD graduation plans. The NSF’s SED logic tree asks doctorates to first select broad definitions of their job types, then to choose the sector, and finally to select or describe work activities. Job types to choose from are limited to six (e.g., postdoc or other training position, employed other than postdoc, further education, etc.). Job sectors to choose from are limited to four – education, government, private or nonprofit, or other, further defined by specific descriptions of the place of work (e.g. “Education”: US 4-year college or university, US medical school, etc., “Government”: US federal government, foreign government, etc.) Graduates are also asked to classify primary and secondary work activities into the following – research and development, teaching, management or administration, professional services, or other.

To summarize, the main career-related categorization tools used in this survey are:

1. Job Type
2. Job Sector
3. Primary and secondary work activities

###### Features

- With annual survey deployment, the SED provides a large dataset for longitudinal comparisons of first-destinations for doctoral degree holders
- Educational history questions allow for longitudinal tracking of the educational path to the doctorate
- Data gathered on financial support shows trends of how doctoral students are supported during graduate school and debt levels related to undergraduate and graduate education
- Broad data fields can be further broken down by factors such as field of study and sex
- Doctoral recipients are surveyed directly

###### Visualization/Reporting

Executive reports are professionally prepared in easy-to read, high-level summaries at regular intervals by NSF. All information is available to download as Excel files or PDFs, and some information is visualized in the prepared reports in bar or line-graph format (https://ncses.nsf.gov/pubs/nsf21308/report/postgraduation-trends#first-postgraduate-position).

##### NSF Survey of Doctorate Recipients (SDR)

###### Basic Definition

The Survey of Doctorate Recipients (SDR) (https://www.nsf.gov/statistics/srvydoctoratework/#qs) is administered every two years and was developed to capture long-term career trajectories of doctoral degree holders from a science, engineering or health field. This survey, conducted by the National Center for Science and Engineering Statistics and the National Institutes of Health, has been conducted biennially since 1973 and is administered to a sample of doctorate recipients from U.S. accredited institutions until they reach the age of 76. Survey data collected via this mechanism focuses more specifically on career pathways taken by science, engineering, and health doctorate holders over time.

###### Development

The SDR collects data on current employment status and occupational information by asking graduates to specify job responsibilities, and their employers’ main business or industry. Employment sectors are categorized further (e.g., self-employed or business owner, private sector employee, U.S. government employee, or other). Educational institution options are surveyed separately, followed by questions regarding the educational institution and academic position. The SDR continues by asking respondents to account for work activities typically engaged in selected from a list (e.g., “Accounting…, Basic Research, Applied Research, “ etc.). Respondents are also asked to categorize their jobs based on a list that is updated periodically. The list of job categories is further divided into specific occupations within each category (e.g. *Job Category*: “Biological/Life Scientist” is broken down into more specific occupations, including “Biochemists and biophysicists,” etc.).

The main categorization tools in this survey are:

1. Employment Sector
2. Work Activities
3. Job Category/Occupation

###### Features

- The NSF’s longstanding history of administering these surveys allows for standardized, longitudinal data collection that enables comparison of trends over time across large, comprehensive data sets.
- The job categories within the SDR are based off of the Standard Occupational Classification system (SOC; the coding scheme for occupations, US Bureau of Labor Statistics, https://www.bls.gov/soc/2018/#classification), thus tying into a robust, tested system that is widely used as a standard for classifying careers.
- The SDR includes granular information about higher education roles (e.g., type of institution, faculty rank, tenure-status, etc.).
- The SDR survey captures over a dozen work activities that occupy at least 10% of the respondent’s time on the job. Additional granular data addresses primary and secondary work activities, type and location of employer, and basic annual salary.
- The taxonomic categories tracked are fairly broad regarding job titles, but multiple functions can be indicated.
- Doctoral degree holders are surveyed directly.

###### Visualization/Reporting

The NSF publishes InfoBriefs on employment among the doctoral scientists and engineers, based on the SDR. All information is available to download as Excel files or PDFs, and some information is visualized in the prepared reports in bar or line-graph format (https://www.nsf.gov/statistics/srvydoctoratework/#tabs-2&rSR&qs&sd&tabs-2&micro&profiles&tools). SDR data are also available to analyze via a special tool termed the ‘Scientists and Engineers Statistical Data System’ (SESTAT). SDR data tables allow for breakdown beyond the major findings in the executive summary and report which focus more on employment status and time to degree (and the intersection with citizenship/international status, gender, etc.), rather than position, title, sector, etc. A multitude of specialized reports analyzing and visualizing various characteristics of the workforce are also available.

##### Council of Graduate Schools (CGS) PhD Career Pathways

###### Basic Definition

The Council of Graduate Schools (CGS) (https://cgsnet.org/understanding-career-pathways) initiated (https://cgsnet.org/join-cgs%E2%80%99s-effort-understand-phd-career-pathways) the PhD Career Pathways project as a multi-phase partnership with a coalition of 75 doctoral institutions and involves collecting information on career outcomes by administering a survey. The CGS Alumni Survey (https://cgsnet.org/understanding-career-pathways) contains questions related to career outcomes, and is inspired by the NSF’s Survey of Doctorate Recipients (SDR) (https://www.nsf.gov/statistics/srvydoctoratework/#qs) taxonomy described above. Broad categorization tools include:

1. Employment sectors (e.g., Education, Government). These sectors are further subdivided based on their characteristics (e.g., Education: research university, liberal arts college; Government: US federal, US state or local, etc.)
2. Job type (e.g., administrator, faculty member, postdoctoral researcher, etc.)
3. Work activities (e.g., managing projects, teaching)

###### Development

The CGS subcategories for educational institutions differ from the 2019 NSF SDR (four-year college or university, medical school, etc.), as the CGS categorizes institutions based on type of institution (e.g., research university, master’s/regional, liberal arts college, community college). Additionally, the CGS’s classification of work activities is based on the NSF SDR, with the SDR 2019 having twice the number of options as CGS. These CGS revisions were made based on the experiences of practitioners using this classification system and their understanding of the shifting career landscape.

###### Features

- The survey asks for information on prior jobs, including secondary paid position(s), which can paint a fuller picture of past and current employment.
- For longitudinal data collection, the survey is administered to three alumni cohorts: those who are 3, 8, or 15 years past their PhD graduation, allowing for career outcome snapshots to be taken at different career stages. The 3-year cohort provides a window on recently graduated PhDs that supplements the NSF Survey of Earned Doctorates (SED) results; the 8-year cohort provides an opportunity for those who entered postdoc positions directly after PhD training to report on their career status; the 15-year cohort allows alumni to share mid-career experiences and any subsequent career changes.
- The survey has evolved since its inception to accommodate participant feedback. As a result, there are several versions of the Alumni and Student Surveys that require institutions to map or crosswalk the data in meaningful ways in order to present and interpret it.
- While the CGS and NSF surveys are similar, comparing results between them can cause challenges because they classify outcomes in different ways

###### Visualization/Reporting

CGS published a series of research briefs (https://cgsnet.org/understanding-career-pathways) based on their analysis of aggregated institutional data. The goal of these briefs is to help campus leaders and analysts contextualize institution-level data, especially in light of the national landscape of PhD career outcomes, while at the same time to continue a conversation about the skills and resources needed for student success in today’s PhD career landscape.

Participating institutions choose how they want to share institutional data. For instance, the University of Wisconsin-Madison has a website dedicated to its participation in the CGS project with information about project goals, data briefs, project highlights, and project timeline (https://grad.wisc.edu/career-pathways/). A majority of data visualizations are created using Tableau or other common data tools that institutions have licensure with, as well as the simple charts enabled by Excel exports (https://cgsnet.org/understanding-career-pathways).

##### First-Destination Survey, National Association of Colleges and Employers (NACE)

###### Basic Definition

The National Association of Colleges and Employers (NACE) (https://www.naceweb.org/job-market/graduate-outcomes/first-destination/class-of-2019/interactive-dashboard/ & https://www.naceweb.org/job-market/graduate-outcomes/first-destination/) aims to provide thought leadership on the relevant issues and trends affecting the college-educated workforce; in doing so, they established national standards and protocols to guide higher education institutions in collecting and disseminating graduate outcomes data. Reporting categories broadly fall into the following: employment status (e.g. employed full time, employed part time, volunteer, seeking employment, seeking further education, etc.); mean and median salaries (full-time employed only); and bonus mean and median. Schools are encouraged to collect other information such as job title, employing organization, and position location, but it is optional to collect this information and these data are not reported to NACE.

###### Development

NACE has collected first destination data on undergraduates for many years. In 2012, they established national standards for NACE member institutions to collect undergraduates’ first-destination outcomes. In 2015, NACE released another set of standards and protocols for collecting information from graduate populations, including both master’s and PhD programs (https://www.naceweb.org/uploadedfiles/pages/advocacy/first-destination-survey-standards-and-protocols-advanced.pdf).

###### Features

- A key benefit of this taxonomy is that it gives NACE-member institutions that were not already collecting graduate program outcomes data a structure to report data.
- This structure aligns with surveys that were already being used for undergraduate outcomes, thus allowing institutions already collecting outcomes of undergraduates to avoid major changes to their survey by applying a similar methodology in order to collect graduate career outcomes.
- The NACE methodology also encourages reporting "knowledge rates”, e.g., reporting the relative percentage of graduates for which an institution has reasonably verifiable information about their outcomes—whether, for example, from self-reported information via surveys, information obtained through public searches (e.g., LinkedIn), or the employers themselves.
- A limitation of this taxonomy is that there is no industry associated with employers, and job titles are self-reported and not standardized with definitions. Job titles are not reported to NACE, though individual schools may report these on their websites. The University of Pennsylvania Career Services reports are an example (https://careerservices.upenn.edu/post-graduate-outcomes/).

###### Visualization/Reporting

NACE reports outcomes both through written reports (https://www.naceweb.org/uploadedfiles/files/2021/publication/free-report/first-destinations-for-the-class-of-2019.pdf) and through an interactive Microsoft Power BI dashboard (https://www.naceweb.org/job-market/graduate-outcomes/first-destination/class-of-2019/interactive-dashboard/) that displays graduate outcomes approximately six months after obtaining their degree. The report can be viewed and/or filtered in many ways, including by degree type (B.S. or M.S.), institution type (i.e., private or public), Carnegie classification type, country, region, Classification of Instructional Program (CIP) code, etc. Prominent within the visualization are salaries and bonuses by career outcome. The outcomes for doctoral degrees are not included in the interactive dashboard but are included in the written report. Furthermore, the report provides the “knowledge rate” mentioned above, as well as the relative percentage of graduates with a known career outcome. The report displays the percent of employed graduates, those that continued their education, individuals seeking employment, graduates who entered the military, and individuals participating in a post-graduate fellowship or internship.

#### INSTITUTION OR CONSORTIUM-DEVELOPED SYSTEMS

##### Unified Career Outcomes Taxonomy (UCOT)

###### Basic Definition

In collaboration with ‘Rescuing Biomedical Research’, the NIH Broadening Experiences in Scientific Training (NIH BEST) Consortium’s doctoral outcomes data was combined with categories used by the Office of Career and Professional Development at the University of California, San Francisco, to yield the three-tiered Unified Career Outcomes Taxonomy (http://rescuingbiomedicalresearch.org/rbr-actions/improving-transparency-ph-d-career-outcomes/).

The UCOT has three classification tiers:

1. *Sector*: These categories describe the broad area of the workforce in which an individual is employed (academia, government, for-profit, nonprofit, and other).
2. *Career Type*: These categories describe the broad type of work performed by an individual within their sector of the workforce. Categories include: primarily research, primarily teaching, science/discipline-related, not related to science/discipline, further training, and unknown.
3. *Job Function*: These categories permit identification of specific skill sets and/or credentials required for employment within that career type. For example, “science writing and communication”, “science education and outreach”, and “science policy and government affairs” are all related to science, yet the function that the individual plays in each role is highly distinct and requires expertise and training that is unique to each function.

###### Development

The NIH BEST consortium curated doctoral outcomes data among Consortium institutions, with the goal of cross-institutional assessment of evidence-based, promising practices for the career development of biomedical PhDs. However, it became clear that the data could not be compared, because each institution curated the data using a variety of different interpretations of the same terms. In an effort to create consistency and reliability in the career outcomes reporting, the NIH BEST Consortium member institutions formed a working group to develop a taxonomy for use within the Consortium^7^. The UCOT provided an initial set of standardized definitions to common terms, which were later empirically tested and clarified to address identified areas of uncertainty. The taxonomy was iteratively tested in Stayart et al 2020^13^ to determine the classification consistency across different ‘raters’; this work resulted in a supplemental guidance document on how to interpret various cases, such that definitions would be applied consistently by practitioners who were curating the data. The results of Stayart et al. (2020)^13^ suggested that reliability improved with all tiers, and improvement occurred even when using non-experienced coders; this experimentally tested, updated version of the UCOT taxonomy was termed UCOT-Exp2.

###### Features

- UCOT is amenable to the addition of customized tracking “flags” for additional granularity that permits further interrogation of the data. This was particularly notable for categorizing faculty appointments, because the flag system permitted the identification of faculty rank and function (e.g., research, teaching, service) and simultaneously identified careers within academia and industry that could be grouped together by common job functions (e.g. leadership, strategy, internal policy, external relations, etc.).
- It has been rigorously and experimentally tested, with a detailed guidance document explaining how to categorize various positions.
- It can be adapted to track those in other disciplines beyond STEM. As an example, Wayne State University has adapted UCOT to the humanities by replacing ‘science-related’ with ‘discipline-related.’ The third tier of the taxonomy can be further adapted by adding additional job functions that are applicable to disciplines outside of STEM^13,14^.

###### Visualization/Reporting

Institutions utilizing UCOT, including institutional members of the Coalition for Next Generation Life Science (CNGLS) (http://nglscoalition.org), report their outcomes data with a wide variety of platforms and visualization methodologies, such as Tableau, static pie charts, and bar graphs. CNGLS members commit to reporting on at least the first two tiers of an earlier iteration of the UCOT taxonomy. A team at the University of California-San Francisco has published a detailed toolkit outlining how they track outcomes using all three tiers of the earlier UCOT iteration^15^.

##### National Institutes of Health Taxonomy (NIEHS-based)

###### Basic Definition

The National Institute of Environmental Health Sciences (NIEHS) developed a three-tiered, hierarchical taxonomy in which postdoctoral fellows are classified by ‘job sector,’ ‘job type,’ and ‘job specifics.’ A detailed description of the taxonomy and how it was developed can be found in Xu et al., 2018^10^ and in the alumni career outcomes dashboard (https://www.niehs.nih.gov/careers/research/fellows/alumni-outcomes/index.cfm).

1. *Job sectors*: describe the broad, overarching areas in which individuals are employed, such as academia, government, nonprofit organizations, for-profit organizations.
2. *Job types:* reflect the relative position levels in which individuals are employed, such as tenure-track positions, non-tenure-track positions, training positions, upper-level management positions, mid-level professional staff positions, and supporting staff roles.
3. *Job specifics:* refer to the duties individuals specifically engage in through their respective positions. Examples of categories within ‘job specifics’ include primarily basic research, primarily teaching, primarily applied research, science writing & communication, and regulatory affairs.

A complete list of *job sectors*, *job types*, and *job specifics* can be found in the Supplemental Tables in Xu et al, 2018^10^.

###### Development

This taxonomy was developed using a “bottom-up” approach, meaning that the career outcomes of NIEHS postdoctoral fellows (who are primarily in the life sciences) were examined, and the designers considered how to best logically bin these career outcomes into categories. The outcomes of nearly 95% of alumni who left NIEHS between 2000-2014 were determined by extensive internet searching and validated by cross-checking with administrative data. Despite developing this with life sciences alumni, the taxonomy has universal applicability for classifying those in both the life sciences and humanities—especially the first two tiers (job sectors and job types). Many of the categories within the third tier (job specifics) are also universally applicable—adopters of this taxonomy could simply add additional categories within the job specifics section to fit their needs (e.g., primarily social science research).

###### Features

1. The three categories (job sectors, job types, and job specifics) are independent of one another.
2. It is unique in that it attempts to codify the relative position level (e.g., management versus support staff, etc.), which helps address questions about under-employment.
3. It is broadly applicable to the sciences and humanities.
4. A benefit to the ‘bottom-up’ approach in developing this taxonomy is that the external labor market guided classification of careers
5. The system contains detailed definitions of each category, as well as example job titles. Additionally, a sample guide is provided that shows how to classify anomymized alumni working for a particular employer, with a given job title, doing a particular type of work activity.
6. It does not capture fine detail regarding faculty-like positions (e.g., adjunct, tenured versus tenure-track). For example, all faculty-like positions are categorized either as tenure-track (which is all-encompassing of tenured, tenure-track, group leader, principal investigator, etc.) or non-tenure-track (research assistant professor, etc.).
7. When using the taxonomy in practice, care should be taken when distinguishing whether to categorize a position as ‘professional staff’ position or ‘management’—management-level classification is typically reserved for those in upper-level leadership positions, often serving in Director-type or Vice-President-type roles.

###### Visualization/Reporting

The postdoctoral career outcomes data are visualized in a variety of ways in order to glean additional insights regarding career outcomes (https://www.niehs.nih.gov/careers/research/fellows/alumni-outcomes/index.cfm). Briefly, the following were used (all based in the R platform): Directional chord diagram, Sankey, Bubble Matrix, Donut, Diverging bar chart, and Geographic visualizations.

##### Track Report and Connect Exchange (TRaCE) (Canada)

###### Basic Definition

Track Report and Connect Exchange (TRaCE) (http://tracephd.com/about-trace/) is a project headquartered at McGill University’s Graduate and Postdoctoral Studies group that aims to track and report on career pathways of PhD graduates, and serves any Canadian institution that would like to partner with them in tracking alumni. The current project tracks humanities, social sciences, and fine arts graduates with both quantitative and qualitative measures with data collection through surveys, data scraping and more recently, narrative interviews. Data collected by surveys and data scraping is used to quantitatively assess overall career outcomes; data collected by narrative interview is reported separately to showcase the stories of how individual alumni navigated their careers (http://tracemcgill.com/wp-content/uploads/2021/03/TRaCE-McGill-QA-Report-Full-Version.pdf)

Career outcomes information collected by survey and data scraping for quantitative analysis includes the following main categories:

1. Employment sector (e.g. academic, government, for-profit, non-profit, etc.),
2. Main field of employer (e.g. education, public and human services, STEM-related, etc.)
3. Job function (e.g. academic research/teaching tenure status, administration, etc.)

###### Development

The current iteration builds on two prior projects: 1) a one-year pilot study in 2015-2016 that tracked humanities graduates; and 2) the TraCE 2.0 project in 2017-2019 that tracked graduates in the humanities, social sciences, and fine arts.

###### Features

- Researchers adhered to a strictly standardized protocol when classifying higher-level data (e.g., sector).
- When categorizing more granular information, such as job functions, the categorization was variable. The research team acknowledged difficulty in categorizing faculty positions and chose to categorize them as non-tenure-track by default if a position’s tenure status could not be verified. While this step may avoid overestimating the number of individuals in a tenure-track-type position, it may have the unintended consequence of underreporting the number of individuals entering tenure-track positions.
- The demographic information collected in the surveys extended beyond basic information and included detailed options for one to self-report their gender identity and sexual orientation.

###### Visualization/Reporting

Data and narratives from 2008-2018 graduating cohorts at McGill University are visualized via an executive summary (http://tracemcgill.com/wp-content/uploads/2021/03/TRaCE_McGill_2103023-1.pdf). Additionally, a quantitative report disaggregates the McGill data in many ways, including a detailed breakdown of the self-identified sexual orientation of participants. (http://tracemcgill.com/wp-content/uploads/2021/03/TRaCE-McGill-QA-Report-Full-Version.pdf).

##### University of British Columbia Career Outcome Survey (UBC) (Canada)

###### Basic Definition

The purpose of the UBC Career Outcome survey (http://outcomes.grad.ubc.ca/) was to systematically determine the career outcomes of its doctoral students who graduated with a PhD between 2005-2013. The UBC taxonomy has two main categories:

1. Employment sector: Higher education, Not for-profit, Private sector, Public sector
  - Higher education professionals are further subclassified (e.g. Research-intensive faculty, Teaching-intensive faculty, postdoc, administrator, term faculty, associate researcher)
2. Job titles: Includes a list of job titles that alumni currently hold. Further details, including a list of the definitions created for each field within the taxonomy, can be found on the UBC website (https://outcomes.grad.ubc.ca/methodology.html).

###### Development

UBC’s Interdisciplinary Graduate Studies Program (IGSP) conducted a pilot survey in 2015 to assess the career outcomes of their graduates. In 2016, UBC extended the project to all of their PhD alumni as described above. The survey was designed to minimize the time required to complete it in order to maximize the number of individuals who would take it. Information on those receiving PhDs in philosophy and English were collected through the national TRaCE project, which, as described previously, collects information on humanties PhDs for partnering Canadian institutions.

The authors used a multi-pronged approach, wherein both surveys and analysis of publicly available data were used to categorize outcomes, of which information was obtained for 91% of graduates. Approximately half the students responded to surveys, and thus information on the remaining students was obtained through internet searches. Survey responses were double-checked according to the alumni’s position and employer, and alumni miscategorizations (relative to UBC’s established taxonomy) were corrected.

###### Features

- UBC compares career outcomes data across various disciplinary groupings by also classifying programs according to the Statistics Canada Classification of Instructional Programs 2000 (https://www.statcan.gc.ca/eng/subjects/standard/cip/2000/index), the categorization system used for sharing U15 university data. This has the advantage of allowing comparisons of outcomes by discipline, rather than only at the individual program level, which is advantageous due to increased group sizes.
- Another strength of this study is that the authors make an effort to comprehensively research each and every position when they are not clear, so as to best characterize it rather than relying solely on either a survey response (which can be miscategorized), or by observing employers and job titles only at a surface level.
- With the current taxonomy, while more detail is available on what individuals are doing within the higher education sector, details are limited on the type or level of work being carried out in other sectors.

###### Visualization/Reporting

The PhD Career outcomes are publicly available online via an interactive dashboard using common visualization types (bar graphs, pie charts, etc.) (https://outcomes.grad.ubc.ca/index.html). The data are disaggregated in several ways, including geographic movement, job location, employers, job titles, gender, domestic versus international, and even down to data source (survey versus internet search). Career outcomes can also be visualized by sector of graduating discipline as well as the specific program of study. A comprehensive report that provides additional visualizations as well as alumni profiles was created and disseminated (https://outcomes.grad.ubc.ca/docs/UBC_PhD_Career_Outcomes_April2017.pdf).

##### University of Toronto-10,000 PhDs Project (Canada)

###### Basic Definition

The purpose of the 10,000 PhDs Project at the University of Toronto (U of T) (https://www.sgs.utoronto.ca/about/explore-our-data/10000-phds-project/) was to determine the current (2016) employment positions of 10,886 individuals who graduated from U of T from 2000 to 2015 in all disciplines.

Career outcomes were categorized in the following manner:

1. Employment sectors: Post-secondary education, private sector, public sector, charitable sector, etc.
2. Within each sector, relative position types or job functions were assigned. Within the post-secondary education sector, position types were defined such as tenure-track professors, full-time teaching stream professors, etc. Within the private sector, functions were defined based on the Government of Canada’s employment categories (such as arts, trades, biotechnology, finance, etc.).

A list of definitions and a detailed guideline for coding are available in Reithmeier et al. 2019 supporting material^16^.

###### Development

The employment positions were obtained by performing internet searches of publicly-available sources such as university, government, company and personal websites, and directories and individual LinkedIn profiles, with ~85% capture success. The School of Graduate Studies (SGS) provided the names of graduates by year and their respective graduate unit/department, gender, immigration status, supervisor and thesis title. No individuals were contacted during the course of this project. Alumni survey instruments were considered but previous studies indicated low returns and the potential for bias based on small sample sizes^17^. Some departments connected with their alumni to create compelling career narratives for their websites.

###### Features

- A strength of the U of T classification system is that it contains granular data on the career outcomes of PhD graduates beyond broad generalizations. The researchers provide a detailed framework that describes each category’s definition—along with an in-depth rationale and logic framework underlying the decision process for classifying individuals in a certain manner^16^.
- The researchers also describe the painstaking lengths to which they went to identify and verify alumni through public sources—listing their commonly used internet sources that provided the most reliable data on alumni career outcomes.
- The U of T researchers also describe how faculty title designations may differ across international barriers, and they provide recommendations for ascertaining the relative equivalencies between Canadian (and U.S.) faculty titles and those from international (non-U.S.) Universities.

###### Visualization/Reporting

The PhD career outcome data is publicly-available on the SGS website using an interactive dashboard (https://www.sgs.utoronto.ca/about/explore-our-data/10000-Ph.D.s-project/). The data can be searched by division, discipline, gender, and immigration status. A 10,000 PhDs Project Overview and Divisional Fact Sheets with clear infographics, created using Tableau, can be downloaded. The 10,000 PhDs Project was initiated as a research project using student researchers with a peer-reviewed publication in PLOS-One as one of the desired outcomes^12^. The U of T has also joined the Coalition for Next Generation Life Science (CNGLS), and they have updated their interactive dashboard to report their career outcomes according to the standards set for those joining CNGLS, which includes reporting via the UCOT 2017 taxonomy format. Reporting data based on these two different taxonomies allows readers to see how similar the Canadian employment sectors are relative to those within the 2017 UCOT (http://rescuingbiomedicalresearch.org/rbr-actions/improving-transparency-ph-d-career-outcomes/), and further updated (UCOT Exp2) based on experimental evidence in Stayart et al. 2020^13^.

#### PROFESSIONAL ASSOCIATION-DEVELOPED SYSTEMS

##### American Association of Universities Data Exchange (AAUDE)

###### Basic Definition

The AAUDE taxonomy (https://www.aau.edu/sites/default/files/AAU-Files/PhD/Project-Summaries-02.22.19.pdf) effort takes advantage of existing government employment classification methodologies, including the jointly developed U.S., Canadian, and Mexican North American Industry Classification System (https://www.census.gov/naics/ NAICS; the coding scheme for industry) and the U.S. Standard Occupational Classification system (https://www.bls.gov/soc/2018/#classification SOC; the coding scheme for occupation, US Bureau of Labor Statistics, 2018). Thus, the AAUDE taxonomy relies on a well-understood, user-friendly coding system where jobs are classified by industry (sector) and occupation (or function). Both industry and occupation have a primary (“major”) and a secondary (“minor”) level, thus enabling fairly specific classifications without being overly detailed and thus burdensome for coders. It also does not require extra “tags” or other designators to code level of work (such as managerial-level or tenure track).

Mandatory fields used to classify alumni in the AAUDE taxonomy are:

1. Top-level employer (Industry) type: e.g. Academic, Industry, Non-profit, Government, Entrepreneurial, Freelance
2. Second-level employer (Industry) type: Institution type or NAICS industry code
3. Top-level occupation: e.g. further study, academic career stage, other research position, other full-time work, exclude from cohort, other (includes non-work occupations such as travel)
4. Second-level occupation: e.g. academic career stage, SOC (occupation code), “other” detail

A full description of the taxonomy is published online (https://www.aau.edu/sites/default/files/AAU-Files/PhD/Project-Summaries-02.22.19-1.pdf).

###### Development

The AAUDE system is based on both NAICS and the SOC, but it sometimes combines and sometimes excludes certain categories or levels of classification, and even adds its own categories for academic careers. The AAUDE classification system was designed to improve data-sharing about PhD career outcomes among AAU institutions. To enable cross-institutional comparisons, a working taxonomy was created in 2017, and then refined in 2018, resulting in Version 6 described above.

###### Features

- A strength of this classification system is that it is sufficiently detailed to capture nearly all career outcomes of PhD alumni across disciplines, but is not so detailed that it would take a coder extensive time to code an employment outcome.
- It is able to reach broadly across disciplines and career pathways, and is thus potentially more useful in coding employment outcomes for humanities students than other existing coding systems. AAU institutions are required to provide their outcomes to AAU annually using this taxonomy.
- The NAICS and SOC codes are standardly used by government reporting agencies, providing robustness and longevity (these codes are available publicly and can also be utilized by trainees for career exploration in O*NET (https://www.onetonline.org/)-the career portal using the federal workforce classification system).
- As a result of the AAUDE taxonomy’s wide use, the data can be compared across institutions and over time. Furthermore, this coding scheme is one of the three being used by the consulting firm Academic Analytics (https://academicanalytics.com/), which is currently being adopted by some universities to gather career outcomes data.
- A limitation of the underlying SOC and NAICS coding schemes is that they are updated infrequently, such that newer career paths may not be represented (created in 1977; last updated in 2018 but revisions can take up to 10 years https://www.bls.gov/soc/2018/#classification).

###### Visualization/Reporting

A large number of institutions collect information as part of the AAUDE initiative. In one such example, Texas A&M University reports the results from their AAUDE Doctoral Exit Survey (https://grad.tamu.edu/Prospective-Students/CNGLS-Coalition-for-Next-Generation-Life-Science/AAUDE-Doctroal-Exit-Survey), which includes interactive drill-down options, enabling one to visualize differences in career outcomes from a variety of groups, including those from different ethnicities.

##### American Historical Association (AHA)

###### Basic Definition

The American Historical Association (https://www.historians.org/wherehistorianswork) serves historians in all professions. As part of serving its constituents, AHA embarked on a project to identify the career outcomes of historians on a national scale. In doing so, it developed a taxonomy to classify their careers. Similar to the AAU taxonomy described above, the AHA taxonomy also includes standard SOC codes from the Bureau of Labor Statistics. The taxonomy collects information in two main categories:

1. Sector: (e.g., government, academia, for/non-profit, etc.). Notably, sectors also include further academic granularity such as definitions for higher ed admin/staff, post-doc, and variants of 2- or 4-year tenure- or non-tenure-track positions; not-found and retired/unemployed are also included in this category)
2. Job function: SOC codes (https://www.bls.gov/soc/2018/#classification) from governmental standardized definitions for job functions

###### Development

Among the vanguard of career outcome transparency for PhD’s, the “Where Historians Work” AHA taxonomy was developed as part of an initiative funded by the Andrew W. Mellon Foundation to track career outcomes for historians nationally and was published as a summary report^18^ and interactive dashboard (https://www.historians.org/wherehistorianswork).

###### Features

- This approach is comprehensive, including all historians graduating with PhD’s nationally between 2004-2013. This list of historians was ascertained by analyzing the names and dissertation titles from the AHA’s Directory of History Dissertations. Using these data, AHA searched publicly available online sources to determine the career outcomes and found data for 93% of historians using the AHA’s Directory of History Dissertations.
- Institutional and personal data (e.g., specialization area, PhD department, geographic area, and gender) were collected, analyzed, and connected with career outcomes data^18^.
- Since this project was intended to serve the needs of humanities (historians specifically), it may not capture some common social sciences or STEM career outcomes (e.g., a category cited as common for historians such as “Library/Museum/Archive” may be less applicable for scientists).
- By using SOC codes, the AHA taxonomy is versatile and can be benchmarked alongside other groups using these same governmental standards. It includes less common job functions that may be applied more broadly by disciplines outside of history.

###### Visualization/Reporting

The results of this study are available to explore on an interactive Tableau dashboard (https://www.historians.org/wherehistorianswork), and include a variety of visualizations, such as tables, geographic locations, and packed bubbles. Data can be filtered down to reveal the name of the PhD-degree granting department, specialization, cohort, or gender, depending on the specific visualization at hand. Swafford and Ruediger (2018)^18^ provide a concise overview of the different stories that can be told by examining AHA’s dashboard.

##### Modern Language Association (MLA)

###### Basic Definition

The Modern Language Association (MLA) (https://www.mla.org/content/download/99761/2283567/Survey-of-PhD-Recipients-2017.pdf) is the professional association for English and Foreign Languages which also embarked on a project to identify the careers of Modern Language PhD’s. Similar to AHA, MLA also received a grant from the Andrew W. Mellon Foundation to collect information relating to PhD graduates. To collect this information, MLA surveyed a random sample of PhD’s. In the survey, respondents were asked to report where they were first employed, and two tiers of outcomes were collected:

1. Job role: (e.g. tenured faculty, tenure-track faculty, administrative, employed outside higher ed, etc.)
2. Job specifics: (varied based on job role: e.g. full time, postdoc fellowship, business, government, non profit, etc.)

In addition to questions relating to their career path since graduating, respondents were asked questions about their job satisfaction and earnings. A full description of the method and findings was published online (https://www.mla.org/content/download/99761/2283567/Survey-of-PhD-Recipients-2017.pdf).

###### Development

The impetus for the survey arose from concerns relating to the shrinking number of full-time tenure-track positions advertised at postsecondary institutions, as well as a desire to learn about the full range of careers pursued by PhD graduates. The report generated from the findings of the 2012 MLA Survey was published in 2015 as “Where Are They Now”, (https://mlaresearch.mla.hcommons.org/2015/02/17/where-are-they-now-occupations-of-1996-2011-phd-recipients-in-2013-2/). In 2017, the MLA contacted individuals from the earlier 2012 survey and invited them to complete a new survey about their employment since they first received their doctorate. While individual survey responses could not be matched to the original survey because they were anonymous—rendering a study that tracked specific changes over time impossible—it is possible to identify overarching career outcome differences in the cohort between two time points.

- This survey aimed to measure the career progress of responders as opposed to only looking at first-destinations post-PhD. It permits a discrete time-based understanding of the career outcome landscape of PhD graduates of English and foreign languages and thus has application in curricular programming to better prepare doctoral students for a range of careers.
- Unlike the federal government’s Survey of Doctorate Recipients (SDR) (https://www.nsf.gov/statistics/srvydoctoratework/#qs), this study is not longitudinal. Survey respondents were anonymous and could not, as a result, be linked to their earlier 2012 survey. This prevents knowing exactly how many respondents left academia and when, as well as other respondents' movement away from one career and into another. Nonetheless, it allows one to observe trends/changes in career outcomes of the overall cohort from one time period to the next.
- The size of the survey is small; of the 1,949 survey respondents for whom email addresses were found, only 310 responded to the survey. Because of this, one should interpret these career outcome results with caution.

###### Visualization/Reporting

For questions relating to type of employment, the report includes a data table as well as pie charts, columns, and cluster columns to render the data easy to read and clear. For all other questions, the report includes pie charts, columns, and cluster columns. All of the visualizations accompany text-based data analysis.

##### Association of Schools and Programs of Public Health (ASPPH) First-Destinations Data Collection

###### Basic Definition

The Association of Schools and Programs of Public Health (ASPPH) is the membership association representing schools and programs of public health which are accredited by the Council on Education in Public Health (CEPH), and includes more than 111 schools and programs which provide bachelor’s, master’s and doctoral (PhD and DrPH) degrees in the public health disciplines.

Beginning in 2014, ASPPH began collecting first-destinations employment outcomes data from member schools and programs, gathered one year post-graduation. Data is gathered by participating schools and programs and reported to ASPPH annually, including:

1. Employment outcome (full-time employed, part-time employed, employed in a fellowship or residency program, continuing study, etc.)
2. Sector of employment (government, academia, for-profit, non-profit, hospital/healthcare) and sub-sector (local government health department, local government not health department, pharmaceutical company, consulting firm etc.)
3. Whether the position is new post-graduation, or a continuation of existing employment
4. Subject of further study, for those pursuing additional education
5. Salary and bonus
6. Degree debt from public health degree

A full description of the methodology and initial findings was published in the American Journal of Public Health; a detailed description of the taxonomy, including definitions, is found within the supplemental appendix^19^.

###### Development

The ASPPH data collection effort was initiated in response to broader needs for data on the public health workforce as well as the outcomes of public health graduates at the bachelor’s, master’s, and doctoral (PhD and DrPH) levels, which were not systematically or consistently captured in the past^20^. The taxonomy was originally designed in 2014 as a pilot project (https://s3.amazonaws.com/aspph-wp-production/app/uploads/2015/07/ASPPH_Graduate_Employment_Pilot_Project_Report_May2015.pdf) to increase enrollment in public health degree programs. It was designed to be used in tandem with reporting for the Council on Education in Public Health, the accrediting body for public health schools and programs, which requires schools/programs to report on employment outcomes but did not have a set standard for collecting data. The “common questions” used in the pilot were formulated with input from ASPPH member schools and programs; the data collection instrument was designed by the ASPPH Data Advisory Committee.

The final collection includes data from a total of 64,592 public health graduates, of whom 53,463 had known outcomes, from four graduating cohort years from 2015-2018. Data was gathered each year, and the number of schools and programs reporting to ASPPH increased from 55 institutions in 2015 to 111 institutions in 2018.

###### Features

- A feature of note is the detail collected within the sub-sectors, which includes those that are particularly relevant to public health. For example, finer detail on government employment (such as “state health department,” “state government, not health department,” “local (county or city) health department,” “local (county or city) government not health department,” “tribal government”), and the for-profit sector (“pharmaceuticals, biotech, or medical device firm,” “health insurance company”) are uniquely positioned to capture details on the public health workforce. Because doctoral graduates in public health are often hired by government agencies, pharmaceutical firms, consulting firms, and so on, this level of detail provides further insight into the connection between these graduates and the public health and healthcare workforce beyond academia.
- Another key feature of this taxonomy is that “Healthcare” is one of the major categorical sectors, rendering it unique (along with TRaCE) amongst the taxonomies described herein. This could be especially beneficial for identifying public health graduates within the healthcare field. However, it would not be as straightforward to compare across other taxonomies who parse healthcare as a subdivision of other major sectors (e.g., University-associated hospitals may be categorized as Academia in other taxonomies; government-associated hospitals (Veteran’s Affairs) may be categorized as the Governmental sector in other taxonomies).
- The survey also gathers data on student loan debt, salary and bonus.
- This taxonomy was developed to look at discipline-specific career outcomes of those with a public health degree. It thus serves as an example of the benefits gained by using a specific taxonomy for specific constituents.

###### Visualization/Reporting

The publication citing these data primarily reports outcomes by degree level and area of study using tables; the results were analyzed primarily with descriptive statistics, with employment outcome status being compared by area of study.

#### SUMMARY OF TAXONOMIES/CLASSIFICATION SYSTEMS

As evidenced by the taxonomies described, a wide variety of methods for classifying the career outcomes of doctoral-degree holders exists. High-level characteristics of these taxonomies include unique developments and applications such as experimental testing and including narratives and skills. First, it is rare to find a taxonomy with experimentally tested reliability and validity (UCOT Exp2). Second, it is notable that one taxonomy combined data with narratives (TRaCE), with the added benefit of being offered comprehensively as part of a national project. Third, another project takes the approach of better understanding career outcomes by identifying the skills and professional development competencies tied to them (CGS PhD Pathways).

In addition to the unique qualities described above, some taxonomies were designed to capture career outcomes in specific fields such as public health (ASPPH), humanities (TRaCE, AHA, & MLA taxonomies), and STEM (such as the NSF SDR); whereas others were designed to be implemented across discplines (AAUDE, UBC, U of T 10,000 PhDs, NACE, and NSF SED). In addition, however, some that were originally developed for STEM fields have been adapted or modified in fields other than their original disciplines (for instance, UCOT and NIEHS were developed for biomedical careers but can be applied across fields) – and yet, a strength of using a discipline-based taxonomy is that it may capture niche careers in that discipline particularly well.

Another commonality across some taxonomies is the reliance upon standardized common labor metrics (NSF SDR and AHA use the Bureau of Labor Statistics SOCs; similarly, the AAUDE relies upon SOC and NIACS). A benefit of taxonomies based in economic standard measures is that it can be more widely applicable and can be compared with other government data such as economic indicators, allowing comparison of career outcomes at institutions to national, regional, and local economic and market trends. Because these standardized classification codes have been developed and vetted carefully, they are widely representative of skills and career areas; however, a potential downside is that these may not be as updated as frequently (e.g., 10-year cycles), and thus they may not always reflect new or emerging fields. In addition, a limitation of the standardized codes is that, while expansive, they may not always be specific enough to accurately capture the variety of career outcomes specific to doctoral alumni. In contrast to using these common labor standards, the NIEHS taxonomy built their system off of the present labor market by first identifying outcomes and employers and then determining how to bin the outcomes into logical categories. The benefit of this approach is that it allows one to flex to a rapidly shifting career landscape, but the outputs cannot be compared to commonly used, robust standards.

Aside from comparing across the taxonomies themselves, one could also consider that the timing of data collected with these systems varies, including some collections which take place prior to, or near, graduation (SED), and those occurring approximately six months past graduation (NACE), one year after graduation (ASPPH), or at longitudinal intervals (SDR). Since it can take time for new graduates to find employment, data gathered shortly after graduation is likely to appear less favorable than that gathered a year or later post-graduation. Additionally, many institutions that are currently collecting outcomes data may capture single snapshots in time, such that the career outcomes they report are reflective of those who left within the past two decades. However one chooses to report, the data should be clearly marked so that individuals examining the data can understand the the time in which a person graduated or left the institution (in the case of postdocs), and the time in which the career outcomes were collected.

Regardless of the system an institution chooses to classify the career outcomes of their alumni, we feel that it is imperative to identify which taxonomy was used, with clear references to the documentation of the taxonomy. This is crucial for individuals examining the data to have an accurate understanding of what the data mean and to ascertain the degree to which data from different departments or institutions can be compared. One suggestion for how to clearly delineate this would be to develop a universal shorthand methodology for tagging all reports and graphics with the taxonomy used. For example, consider the UCOT taxonomy—when the taxonomy was revised based on feedback and tests to ensure inter-rater reliability, a new version was published^13^. The original version is referred to as UCOT 2017, and the updated version UCOT-Experimental (UCOT-Exp2). This helps to clearly distinguish the specific taxonomy and version being used, which is important, given that taxonomy development should be viewed as a continuing process as career paths shrink and grow throughout our ever-changing economy.

To complement the description of the taxonomies described above, we collated and aligned the major taxonomic categories from a number of additional Universities and classification systems, including all of the taxonomies described herein to create a crosswalk tool (https://osf.io/dwnrk/).^11^ From these alignments, clear patterns and commonalities across classification systems are apparent, resulting in the emergence of an overarching primary list of terms and definitions. For example, all systems include a) **employment sector** in some form, and most also include b) **position/job or career type,** and/or c) **function/role/work activity**.

Within the **employment sector**, eight clear categories appeared most frequently, including variations on the following (number of times sector included): Academic (30X), Government (30X), For-Profit (26X), Non-Profit (28X), Individual (27X), Unknown (21X), Not in workforce (23X), and Unemployed/Seeking (11X). Healthcare was represented as a standalone subsector in two taxonomies, but most taxonomies included healthcare systems within the other 8 sectors described (e.g., University-affilated hospitals would fall under the academic sector while Veterans’ Affairs hospitals would fall under government, etc.). The main area of discrepancy in the sectors described included whether an entity was public or private, and how that was categorized across different taxonomies. For example, Universities could fall under either the public or private sector, as could primary/secondary schools. However, some taxonomies categorized only those at Universities within the academic sector, whereas those in primary or secondary schools were categorized according to whether they were either public/government or private/non-profit. Other taxonomies, on the other hand, categorized all educational institutions (whether preschool through Universities) within the academic sector.

For the remaining two themes (**position/job or career type** and **function/role/work activity**), classification across the taxonomies was less consistent, but broad commonalities could be ascertained. Within position or career types, Faculty-like positions (encompassing those including tenured, tenure-track, tenure-unclear, and group/team leader) were the single most commonly observed category, appearing 45 times in some form. Other commonly observed categories were that of Non-tenure track (27X), Mid-level Professionals (13X), Senior Management (12X), and Trainee (35X). Less common, but appearing multiple times, were Unknown (3X), Discipline-related (6X), Non discipline-related (3X), and Support staff (3X).

For job functions/roles/work activities, 31 categories were able to be aligned among the taxonomies examined, with some appearing at a much higher frequency than others. The full listing can be found in the crosswalk table (https://osf.io/dwnrk/)^11^. Some of the most commonly appearing categories include those conducting research or teaching. These are often further subdivided in order to better ascertain the type of work being conducted (e.g. basic or applied research; full-time faculty teaching or science education/outreach, etc.). Categories most difficult to align concerned those within product development and manufacturing/engineering, as the design and development of products may involve engineering principles. Entrepreneurship was also difficult to align because it is sometimes considered a job function, while other taxonomies include this within the ‘sector’ fields as the ‘Individual’ sector.

The goal of collating and describing the taxonomic classifications for doctoral-level career outcomes is to assist institutions in determining the taxonomy that works best for their needs by highlighting key features, benefits, and caveats to different systems. There is little cost to choosing a taxonomy, but there is a high cost to *not* collecting outcomes data. This cost of not collecting outcomes may become evident in many ways—by obfuscating where students or postdocs enter into careers; in delayed curricular innovations; in difficulties during recruitment by not being able to speak on graduates’ outcomes; and in the cost of an institution not knowing how their doctoral graduates and postdoctoral scholars are contributing to innovations within the global economy and society as a whole. In short, there are many valuable taxonomies from which to choose—the truly costly choice would be to not select any system of reporting the graduate outcomes.

### 2. DATA VISUALIZATION

#### Introduction

In addition to identifying career outcomes and classifying them according to a taxonomy, it is important to communicate these data in an effective manner. In the age of big data, a wide range of visualization methodologies and platforms have become available that can be leveraged for identifying and sharing career outcomes trends. Different visualization techniques can be used depending on the intended purpose and audience. For example, do you want to tell the story of how students from different programs have significantly different outcomes? Do you want to tell the story of how career outcomes have changed over time? Do you want to tell the story of how individuals have migrated from training locations to employment locations? Do you want to tell the story about how individuals from underrepresented backgrounds have career outcomes that fare differently than those from well-represented bakgrounds? The answers to these types of questions can help determine the best course of action when choosing ways to visualize data. Several resources exist to help inform the decision-making process around which visualization method to use: (https://www.klipfolio.com/resources/articles/what-is-data-visualization), (https://depictdatastudio.com/introducing-the-essentials/), (https://github.com/ft-interactive/chart-doctor/blob/master/visual-vocabulary/poster.png), (https://extremepresentation.typepad.com/files/choosing-a-good-chart-09.pdf).

Apart from telling a story, it is also worth considering the way data are visualized so that the data can be comparable for benchmarking purposes and to reduce the likelihood of misinterpretation. As an example, consider whether or not the ‘unknowns’ are included within the data set being visualized. If they are excluded, then the career outcomes values are artificially inflated relative to the true population, since the denominator is artificially smaller by excluding unknowns. As another example, if one were to visualize a subpopulation within an overall student alumni cohort and represent that subpopulation on a scale of 0-100%, a casual reader could easily misinterpret this as representative of the total population, especially if the data were shown out of context. Thus, care should be taken to ensure that visualized data are clearly labeled in all cases. As a way to assess labeling clarity, assume the figure or visualization of the data will stand alone—if taken out of context in this manner, could it be easily misinterpreted? If the answer is ‘yes,’ then the author should either label the figure more clearly or represent the data in a different manner altogether. This is increasingly relevant in the age of social media when snippets, excerpts, or visualizations are commonly highlighted out of context.

##### Visualization Platforms

Below, we list platforms/software that are widely-used to display career outcomes data and stories.

##### Tableau

Tableau software supports popular visualizations such as tables, charts, maps, time series, etc. Despite its simplicity, Tableau is a powerful tool with advantages including integrating querying, exploration, and visualization of data into a single process^21^. It is shown to have high performance with large data sets and can connect to a varied set of data sources^22^. Tableau is capable of providing *ad-hoc* analyses and has provisions to analyze data offline. A potential limiting feature is that Tableau is constrained to generating visual presentation grids with uniform and limited granularity and dimensionality^23^. Nonetheless, when representing outcomes data, this uniformity may be desirable if one wants to compare outcomes between institutions.

Some selected examples of using Tableau to visualize career outcomes data at various institutions include Stanford University (https://tableau.stanford.edu/t/IRDS/views/StanfordPhDAlumniEmployment/StanfordPhDAlumniEmploymentDashboard?:embed_code_version=3&:embed=y&:loadOrderID=0&:display_spinner=no&:display_count=n&:showVizHome=n&:origin=viz_share_link), Johns Hopkins University (https://oir.jhu.edu/phd-career-outcomes/), the University of California-San Francisco (https://graduate.ucsf.edu/program-statistics), and the University of North Carolina at Chapel Hill (https://bbsp.unc.edu/professional-development/career-outcomes/). Institutions vary in how they visualize data within Tableau, yet a significant proportion tend to display data in bar charts and tables, with dropdown filters available to further explore and parse the data.

##### NIEHS custom-made R platform

NIEHS built a custom data platform (https://www.niehs.nih.gov/careers/research/fellows/alumni-outcomes/index.cfm) using open source R language (https://www.r-project.org/), RStudio (https://www.rstudio.com/), and the R Shiny (https://shiny.rstudio.com/) package that allows users to build interactive web apps and dashboards straight from R. A software add-on known as Plotly (https://plotly.com/r/getting-started/), an R package for creating interactive web-based graphs via the open source JavaScript graphing library, was also incorporated into the interface to increase the level of interactivity with the graphics. The platform functions by reading an Excel file and presenting data in a standardized output format. An advantage of using this platform is that it provides full control of how data are presented—including in the nature and types of graphics shown. The platform also allows users to easily present binned data (e.g., 5-year bins, or any desired bin size), which can allow one to avoid presenting data with small sample sizes—thus addressing potential privacy concerns while also making data interpretation more robust. The graphing possibilities using R are plentiful: basic diverging bar charts; donut charts; point-range charts; box-and-whisker plots; Sankey graphs; bubble plots overlaid with heat maps; directional chord diagrams, etc. These visualizations could showcase: a) how demographics change over time; b) how each of three taxonomic tiers relate to each other; c) how country of origin relates to location of job employment; d) how training times differ over time and according to career outcome; e) how outcomes differ as a function of country of origin while simultaneously overlaying either gender differences, training time differences, job location differences, and more.

To date, NIEHS has adapted this platform for several universities’ internal use, and has included key additions such as whether an individual was on a T32 training grant or received their own funding. In order for another institution to use the platform, they need only to format their data in a pre-specified manner (Download example “Source Data” file located at the bottom of the NIEHS Alumni Outcomes Dashboard: https://www.niehs.nih.gov/careers/research/fellows/alumni-outcomes/index.cfm). The R platform can then read the Excel file and instantly produce the dashboard as an output. If an institution’s data are not formatted in the manner shown, they can collaborate with NIEHS to adapt the platform to their needs or work within their own institution to modify the R-code for their own purposes. The code, as well as a comprehensive handbook (https://github.com/nihxuh/alumni-customization/commit/17451228e31c38c8d56d9b85f54699428cae1e54) containing instructions on how to use and modify the code, can be found for both the original dashboard version (https://github.com/nihxuh/alumni-shiny) and for the updated version (https://github.com/NIEHS/alumni-dashboard/).

##### Microsoft Excel

MS Excel is a ubiquitous, low-cost, and easy-to-use tool typically used for basic data analytics. It affords a first step to data visualization, with multiple types of first-order charts. Basic charts available include column, line, pie, bar, area, and scatter plots, and for more sophisticated visualizations, stock, surface, bubble, donut and radar charts are available. Some variability exists within each chart type, such as 2-dimensional vs. 3-dimensional visualization, use of color, etc. Excel visualizations for PhD career outcomes data include divergent stacked bars, word clouds, and Sankey diagrams (e.g., Oregon Health & Sciences University, https://www.ohsu.edu/sites/default/files/2020-04/OHSU%20SoM%20Outcomes%202019%20Report%2004082020.pdf; for a tutorial, see: https://peltiertech.com/diverging-stacked-bar-charts/). There are a number of training resources for improving Excel visualization skills; one such example can be found here (Depict Data Studio: https://depictdatastudio.com/visualizing-equity-in-education/). For those desiring more advanced graphic functionalities, a Poweruser plugin (https://www.powerusersoftwares.com) can be installed that will enable one to create more complex graphics within Excel, such as Sankey diagrams.

##### Career Service Management (CSM) Platforms

Three of the most popular CSMs on the market are Symplicity (https://www.symplicity.com/), Handshake (https://joinhandshake.com/career-centers/?_ga=2.134659441.1874268023.1607558995-968462549.1607558995), and 12Twenty (https://www.12twenty.com/)—each of which offers modules that permit institutions to track, measure, and visually present their first destination survey outcomes. Symplicity, Handshake, and 12Twenty allow institutions to develop and launch first destination surveys tailored to their needs and, in the case of Symplicity, integrate the NACE First Destination Survey (FDS) into their CSM dashboard. These are user-friendly systems that are easy to navigate—especially for those who lack data skills, as the data analysis is automated by the CSM.

The integration of first destination outcomes into CSMs is relatively recent and reflects the growing interest of educational institutions to provide statistical evidence of their students’ success as well as to potentially provide prospective students with data pertaining to their programs to help them make informed choices. While the data collected through the CSM are frequently presented on outward-facing webpages, the integration of first destination survey outcomes in the CSM permits current students and alumni to identify career paths, industry trends, and alumni employment locations associated with their respective programs.

First Destination Survey questions may be integrated into CSM, or data collected through surveys managed outside of the CSM may be uploaded into them following the CSM-designated method. Users (students and administrators alike) can select from predetermined and/or customized fields to view visualizations of the data. FDS outcomes may be included as part of a platform or as add-ons depending on the particular CSM.

##### Microsoft PowerBI

Power BI (https://powerbi.microsoft.com/en-us/) is a business intelligence (BI) analytics service by Microsoft with both cloud-and desktop-based interfaces. Data can be imported into Power BI from a variety of sources such as Microsoft Excel, MailChimp, Salesforce, etc. through a breadth of data connectors. Large datasets, commonly referred to as “Big Data”, can also be integrated directly into the Power BI web service, which allows for easy sharing. Power BI boasts an intuitive user interface. Once data are loaded, users can click through a variety of options to choose the desired visualization, which includes Sankey diagrams, bullet charts, aster plots, word clouds, and more. Formats for various visualizations can be easily customized, and many options are also available for interactive visualizations. More complex tasks such as joining datasets can also be easily accomplished. With both the simplicity in building visualizations and the plentiful customization options, Microsoft Power BI is generally a highly rated tool. Some selected examples of graduate-level career outcomes showcased with Microsoft Power BI include Wayne State University (https://oira.wayne.edu/dashboard/graduate-school/phd-alumni-survey-report), the University of Texas System (https://seekut.utsystem.edu/GradNat), and Weill Cornell Medicine (https://mdphd.weill.cornell.edu/alumni-and-outcomes/career-paths-our-graduates). NACE, as mentioned above in the taxonomies, showcases the collective outcomes of college undergraduates using Power BI (https://www.naceweb.org/job-market/graduate-outcomes/first-destination/class-of-2019/interactive-dashboard/). Power BI Desktop is available to download for free, while other options (e.g., Power BI Pro) have subscription costs, and include features such as API embedding, peer-to-peer sharing, and support for data analysis. One limitation to note is that Power BI only functions within a Microsoft Windows environment; individuals with other operating systems must utilize a virtual Windows desktop to run Power BI.

##### Prism, IBM SPSS, & SAS

Some data analysis software packages are also equipped to provide accompanying visualizations using check-box and pre-programmed options. This is especially useful for individuals without coding experience. Both Prism (https://www.graphpad.com/scientific-software/prism/) and SPSS (https://www.ibm.com/products/spss-statistics) are examples of plug-and-play software platforms that allow the use of default settings and that help guide users in choosing which statistical tests to use based on the data selected. Prism, which is oriented toward life-science examples, is especially user-friendly with tutorials and sample data sets, whereas SPSS is oriented toward complex analyses using control variables and was originally developed to support research in the social sciences. Both software packages produce useful visualizations such as graphs, box-and-whiskers plots, scatter plots, and more. SPSS visualizations are typically useful while completing analyses, but other software packages produce more customizable variations reproduced for publication-quality visualizations (e.g., R, IBM SAS; https://www.sas.com/en_us/home.html, Prism, or other customizable visualization tools). Prism is easily customized and can create publication-quality visualizations, though some of its advanced analytics are less developed than the default options available in IBM SPSS. SAS provides innovative visualizations and robust data storage abilities, and it supports customized complex analyses. However, SAS requires coding knowledge to fully use these features. Each of these software options can provide basic analyses and basic visualization tools, so available institutional training and support may dictate adoption at the department, office, or institutional level.

##### Institution-Specific (Customized Platforms)

Some institutions have developed homegrown solutions that are fully customized for their needs. This may be a viable option if internal institutional resources are available to assist, if the staff member responsible has skills in this area already, or if trainees at the institution can be trained to support development of the customized system. Alternatively, institutional or grant funds may be leveraged for an initial one-time investment to set up a complex customized system if its maintenance over time is possible long-term or if an ongoing contract is feasible. For instance, UBC used a Java-based (JQuery) hosted on a Google library (https://outcomes.grad.ubc.ca/comp-cohorts.html & https://developers.google.com/speed/libraries#jquery), and Clemson University similarly used a java-based Google library (https://career.sites.clemson.edu/data_analytics/FDS_App.php?year=18_19&filters=%60degree%60=%22Doctoral%20degree%22#open) while Boston University used a Plotly Javascript library (https://www.bu.edu/grad/Ph.D.-profiles/SPH-Epidemiology_Profile.html). While these examples are not intended to be comprehensive, they illustrate some examples of institutions that have developed their own fully customized platforms to display their data.

###### Visualization Types

As with the other sections of this report, the following list of visualizations is intended to be informative but not exhaustive. While outcomes data can be visualized using basic pie charts, donut charts, bar charts, and line charts, we have chosen to include the career outcomes visualization examples that showcase data in innovative ways. Resources for deciding which visualization can best illustrate your career outcome stories can be found here (https://depictdatastudio.com/charts/).

##### Diverging Stacked Bars

Diverging Stacked Bars can allow for visualization of changes over time, such as with career trajectory or demographics. As an example, snapshots of data for the graduate population at Oregon Health & Sciences University (OHSU; https://www.ohsu.edu/sites/default/files/2020-04/OHSU%20SoM%20Outcomes%202019%20Report%2004082020.pdf) are captured at specific time points after completing their PhD (e.g., 1st year, 5th year, 10th year) (Fig. 1). For each Year-Post-Graduation cohort, the graduates are represented as a bar with various colored segments—each color of which depicts the relative percentage of alumni in a particular career type, with the total of segments equaling 100% of the cohort. These cohort years are stacked on top of each other, and a divergence point is chosen. In this case, OHSU chose the divergence point as postdoctorates and tenure-track faculty. When each cohort is stacked one on top of the other, it becomes immediately apparent when the proportion in the largest career type (in this case, postdoctorates) changes across the different cohorts based on how many years have passed post-graduation. As the number of years postgraduation increases, the number of alumni in postdoctorate positions decreases, while the number in tenure-track faculty increases. Increases in other careers can also be seen, though the largest movement appears to be in the transition from postdoc to faculty. As indicated in the graphic, the raw number of PhD graduates surveyed across each cohort differs; it is important to note these differences when visualizing data so that readers can ascertain the limitations of data interpretation. In another example, diverging stacked bars are used to show how population demographics of trainees at the NIEHS^10^ changed over time, from a higher percentage of males (over 60%) in the early 2000’s to a more balanced, nearly 50/50 population of males and females in the 2010’s. These types of graphics can be created in a number of programs such as Excel (e.g., see http://stephanieevergreen.com/diverging-stacked-bars/) and are also readily made within other software and programming languages (including Tableau, Python, R, etc., see https://towardsdatascience.com/diverging-bars-why-how-3e3ecc066dce).

**Figure 1:**
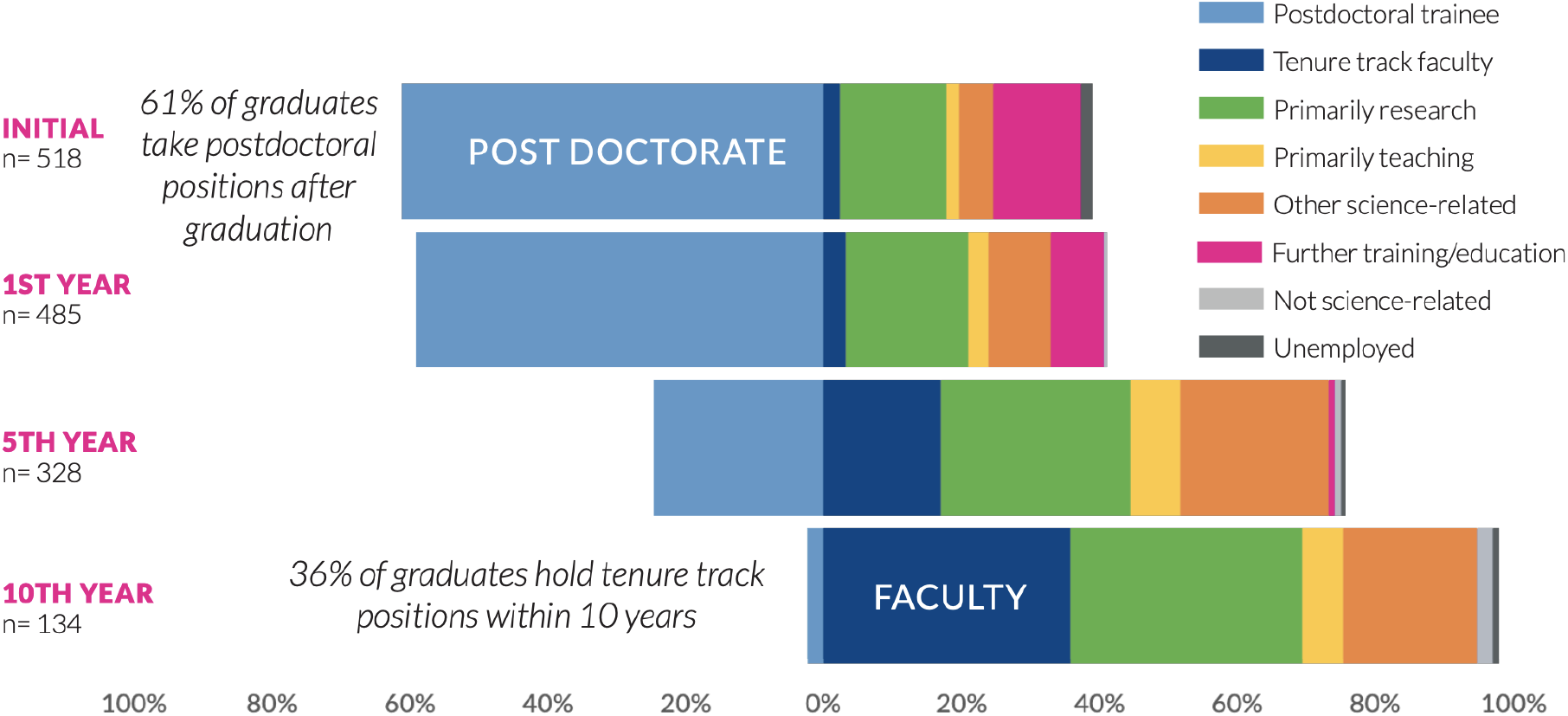
Using Diverging Stacked Bars to Visualize Changes in Career Outcomes Over Time. Diverging stacked bars showcase the relative percentage of PhD graduates from the Oregon Health & Science University School of Medicine entering into careers at given points after graduation (initial position; 1 year after graduation; 5 years after graduation; 10 years after graduation). 36% of graduates become tenure-track faculty within 10 years after graduating. Figure provided by Allison D Fryer, Jackie Wirz and Amanda Mather, Graduate Studies School of Medicine, Oregon Health & Science University. Unpublished data. Reproduced with permission, available at: https://www.ohsu.edu/sites/default/files/2020-04/OHSU%20SoM%20Outcomes%202019%20Report%2004082020.pdf)

##### Sankey Diagram

A Sankey diagram allows for complex visualization of the relationships between two or more variables with multiple possible outcomes. For example, a Sankey diagram can show how individuals with degrees in different fields ‘flow’ into different careers upon graduation – examples include diagrams from Wayne State University^24^, Stanford University (https://irds.stanford.edu/data-findings/phd-jobs), and Purdue University^25^. This same type of flow has also been shown in terms of geography, in which alumni from a given country ‘flow’ into a career either in the same country or a different country (e.g., UBC; https://outcomes.grad.ubc.ca/geographic-movement.html).

Furthermore, a Sankey diagram can be used to emphasize relative proportions; one example from the National Institute of Environmental Health Sciences^10^ visualizes three tiers of a taxonomy (Fig. 2). Taking the academic job sector (shown in red) on the far left as an example—as the viewer moves to the right visually, career outcomes in the academic sector are further divided into professional, management, tenure-track, support, non-tenure-track, and trainee job types. If one focuses on the tenure-track job type (shown in green) and continues moving to the right, one can see that the main proportion of those in this job type are conducting basic research. An interactive form of this diagram is also available (https://www.niehs.nih.gov/careers/research/fellows/alumni-outcomes/index.cfm; see ‘Relationship between categories’). A potential pitfall of using a Sankey diagram to illustrate proportions is that the data may be misinterpreted as ‘flow,’ meaning, for example, that someone may think an individual is moving from an academic career to a management career to a basic research career. Therefore, it is critical to clearly label the Sankey diagram when it is used in this manner. A Sankey diagram was chosen in Fig. 2 because it clearly and effectively illustrates how the career outcomes from all three tiers of a taxonomy are related to each other, which is not possible when each tier is represented separately.

**Figure 2.**
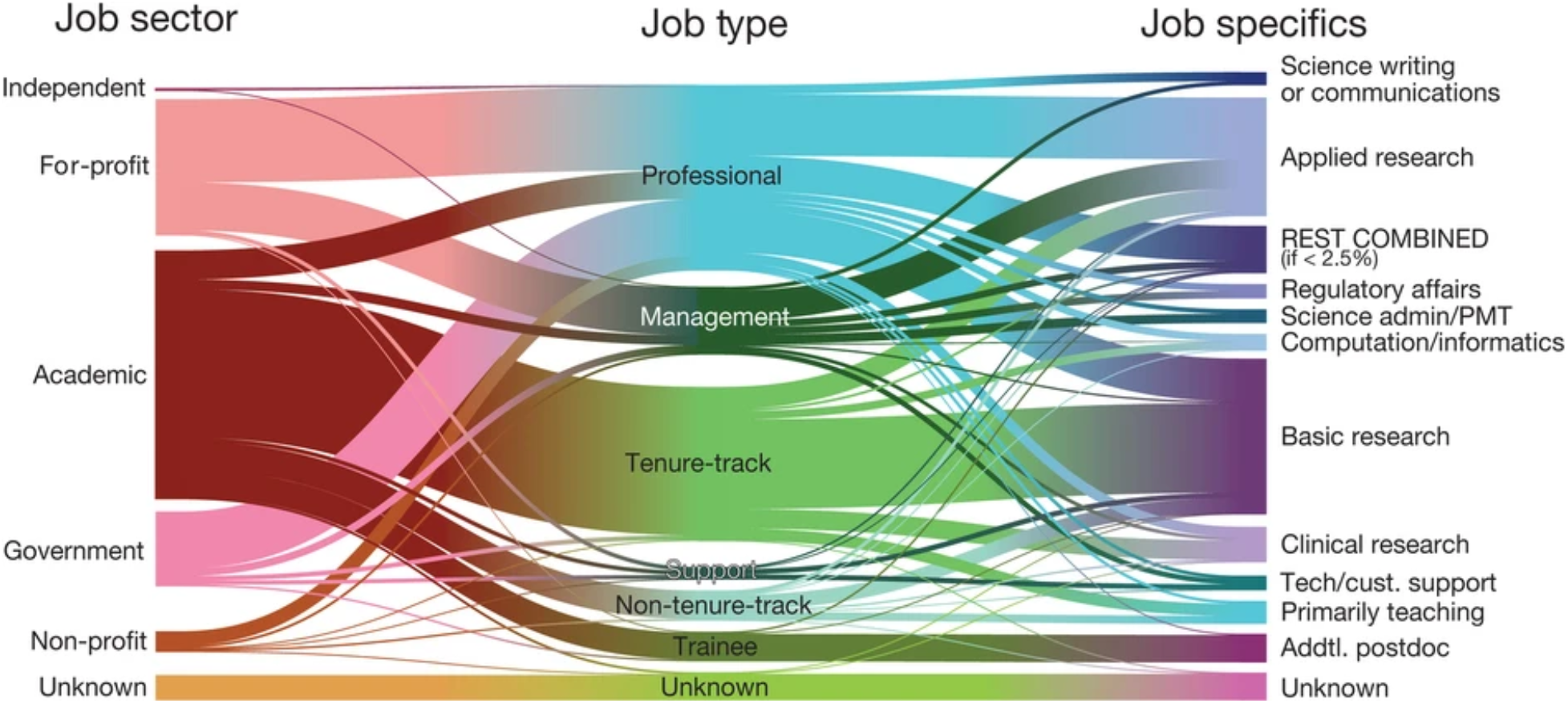
Using a Sankey Diagram to Illustrate Career Outcomes and the Relationship Between Tiers of a Taxonomy. A Sankey diagram shows the relationship between three tiers of a taxonomy from postdoctoral alumni at the National Institute of Environmental Health Sciences. Moving from left to right across the diagram, one can see, for example, that the majority of those who enter academia do so as tenure-track faculty. Conversely, it is apparent that most who enter tenure-track faculty are in academia, with a few in the government sector and even fewer in the non-profit sector. Continuing from the middle and moving to the right, it is clear that the majority of those in tenure-track faculty positions are conducting basic research, while a smaller proportion are in applied research, clinical research, or teaching positions. Reading from right to left, it is apparent that most individuals conducting basic research are in tenure-track positions, with a smaller proportion in other types of positions. (Reproduced without changes; https://creativecommons.org/licenses/by/4.0/)^10^

Sankey diagrams can be made in R or Python (https://www.data-to-viz.com/graph/sankey.html) as well as in many of the platforms described above such as Tableau and Microsoft Power BI. They can also be created in Excel if the add-in power-user (https://www.powerusersoftwares.com/) is installed. Sankeymatic (http://sankeymatic.com/) is also a helpful tool for creating a Sankey diagram that does not require coding experience or any additional external software. Users input the data into the online interface, and export the diagram as a jpeg, which can be further formatted using other software such as Adobe Illustrator or Photoshop.

##### Directional Chord

A directional chord diagram is similar to the Sankey in that it allows one to analyze the flow between different sets of entities. The entities are displayed around a circle and are connected by arcs. What sets this apart from a typical chord diagram is that it provides directionality, with arrows pointing in the direction of flow, thus making it more apparent to the viewer. In an example from NIEHS, we can see the flow from country of origin into country of employment where arrows point to employment location^10^ (Fig. 3). Examining Japan, for instance, one can see that nearly all fellows with Japan as their country of origin return to Japan for employment (note the thicker orange arrow/arc). Conversely, a much smaller proportion of fellows from Japan obtain employment within the USA (note the thin orange arrow/arc). At the same time, it is also apparent that there is little flow into employment in Japan from those who originate from countries outside of Japan. In contrast, if we examine China, one can see a thicker purple arrow denoting flow into employment within the USA, with a thinner arrow returning to China. Similar to Japan, there is little flow into employment in China from those with a different country of origin.

**Figure 3.**
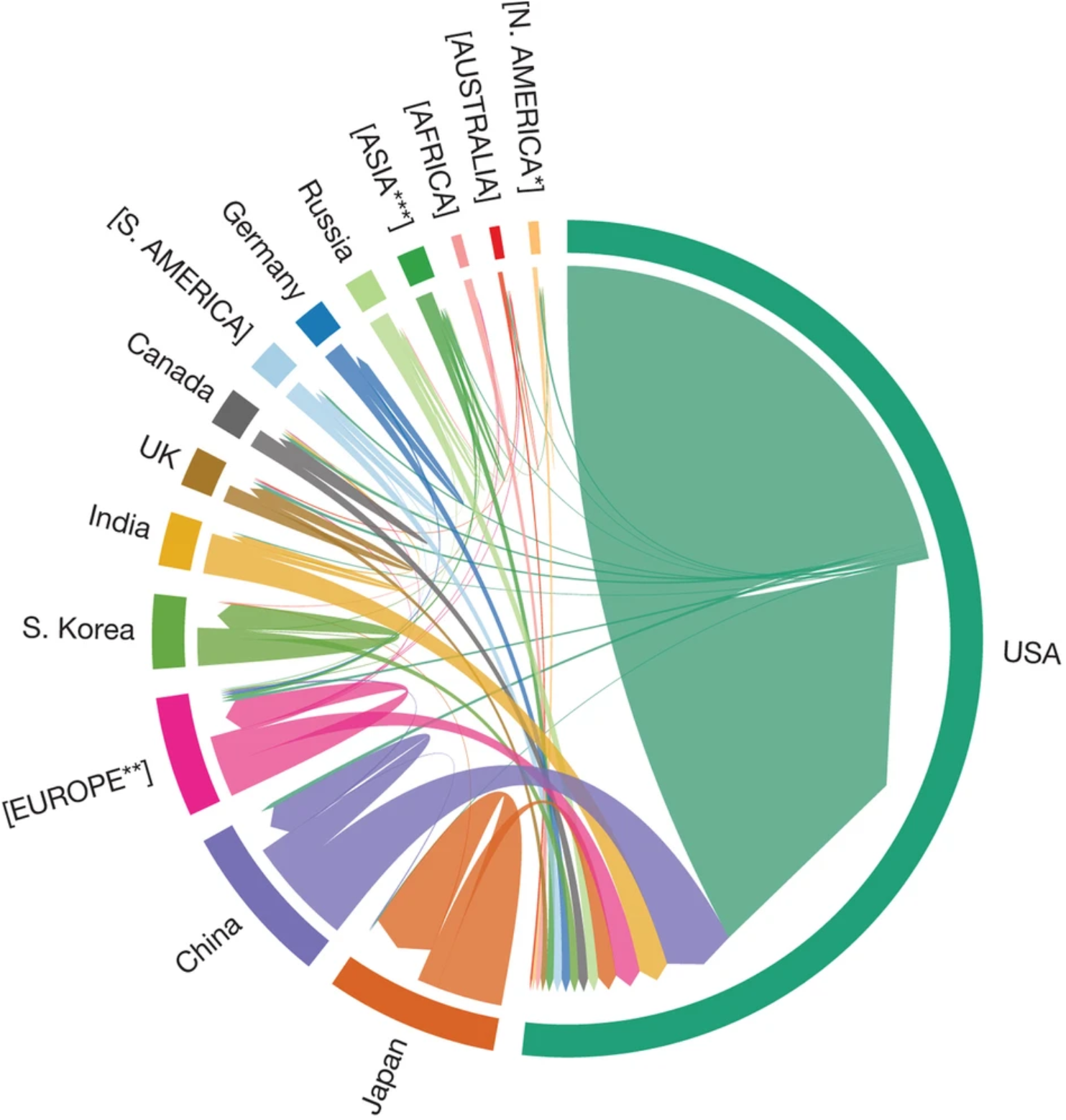
Using a Directional Chord Diagram to Illustrate the Relationship Between Country of Origin and Country of Employment. This directional chord diagram displays the countries of origin alongside the countries of job location (arrows point to job locations) from postdoctoral alumni at the National Institute of Environmental Health Sciences. Nearly two-thirds of alumni remain in the US after training; fellows from Japan, South Korea, the UK and Germany enter into careers in their home countries more so than fellows from other countries. *North American countries excluding US and Canada; **European countries excluding UK, Germany; ***Asian countries excluding China, Japan, India and South Korea. If there are enough alumni to visualize from an individual country, it is shown in title case. Remaining countries are grouped and depicted by continent in all caps. (Reproduced without changes; https://creativecommons.org/licenses/by/4.0/)^10^

##### Stacked Donut or Sunburst

A stacked donut provides a simple way to visualize a snapshot of outcomes, using, for example, the UCOT 2017 three tier taxonomy deployed by Wayne State^26^ (Fig. 4). The stacked donut typically consists of concentric circles; in this example, three circles are stacked, with each representing a different point in time from when graduates received their doctoral degrees. The inner circles represent those 0-5 years from receiving their degree; the middle circles represent those 6-10 years out, and the outermost circles represent those 11-15 years from receiving their degree. Career outcomes for each tier of the UCOT 2017 taxonomy are shown, with the top plot representing the employment sector (Tier 1), the middle plot representing the career type (Tier 2), and the bottom plot representing job function (Tier 3). Upon examining the plots, it is readily apparent that the proportion of individuals engaged in further education or training (see Tier 2) declines sharply the further out one is from receiving their degree; likewise, the proportion entering into either faculty or group leader positions (see Tier 3) significantly increases the further out one is from receiving their doctoral degree.

**Figure 4.**
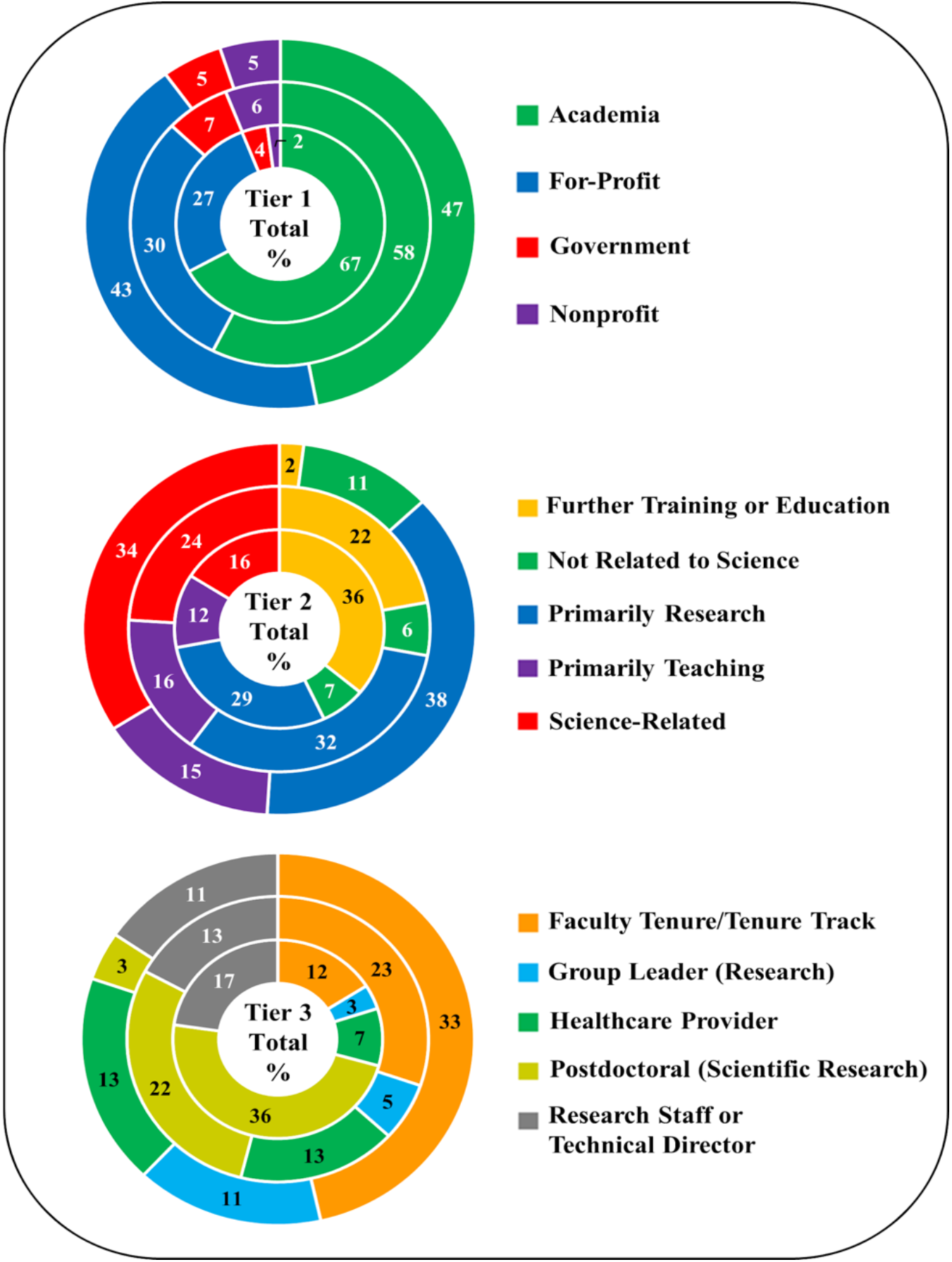
Using Stacked Donut Plots to Illustrate Career Outcomes from Three Taxonomic Tiers at Different Points in Time. These stacked donut plots depict the career outcomes of Wayne State University’s biomedical doctoral alumni at different points in time after receiving their degree (either 0–5 years, inner circle; 6–10 years, middle circle; or 11–15 years, outer circle). The top plot represents outcomes of alumni by employment sector (tier 1); the middle plot represents outcomes by career type (tier 2), and the bottom plot represents outcomes by job function (tier 3). (Reproduced without changes; https://creativecommons.org/licenses/by/4.0/)^26^

##### Bubble Plot & Heatmap

A bubble diagram is a powerful visualization tool for multiple variables simultaneously. When overlaid with a heatmap, one can add even greater dimensionality to the data. As shown in an example from NIEHS, career outcomes are separated by country of origin, job type (and relative percentage within that job type), as well as time in postdoctoral position^10^ (Fig. 5). If we compare the professional staff positions as an example, it is apparent that the greatest proportion of U.S. fellows enter into these types of positions. Examining mean training time shows that they spend on average between 30-40 months in training. If we compare this to fellows from Japan, stark differences emerge—for example, it is clear that fewer fellows enter into professional staff positions relative to the population of fellows from Japan. At the same time, it is also clear that the training time is significantly greater (approaching 60 months) for those fellows from Japan who enter into professional staff positions. While a lot of stories can be told with this type of visualization, one limitation is that it is difficult to discern differences in bubble size, thus restricting one’s ability to accurately estimate true percentages. Furthermore, the high degree of data dimensionality could confuse the casual reader. Figure 6 shows another example of a bubble plot reproduced with permission from the American Historical Association’s project to identify where historians work (https://www.historians.org/wherehistorianswork). This plot is more straightforward in that it depicts the relative proportion of alumni entering into careers classified by SOC codes. An interactive version of this plot can be found found on the AHA website (https://www.historians.org/wherehistorianswork).

**Figure 5.**
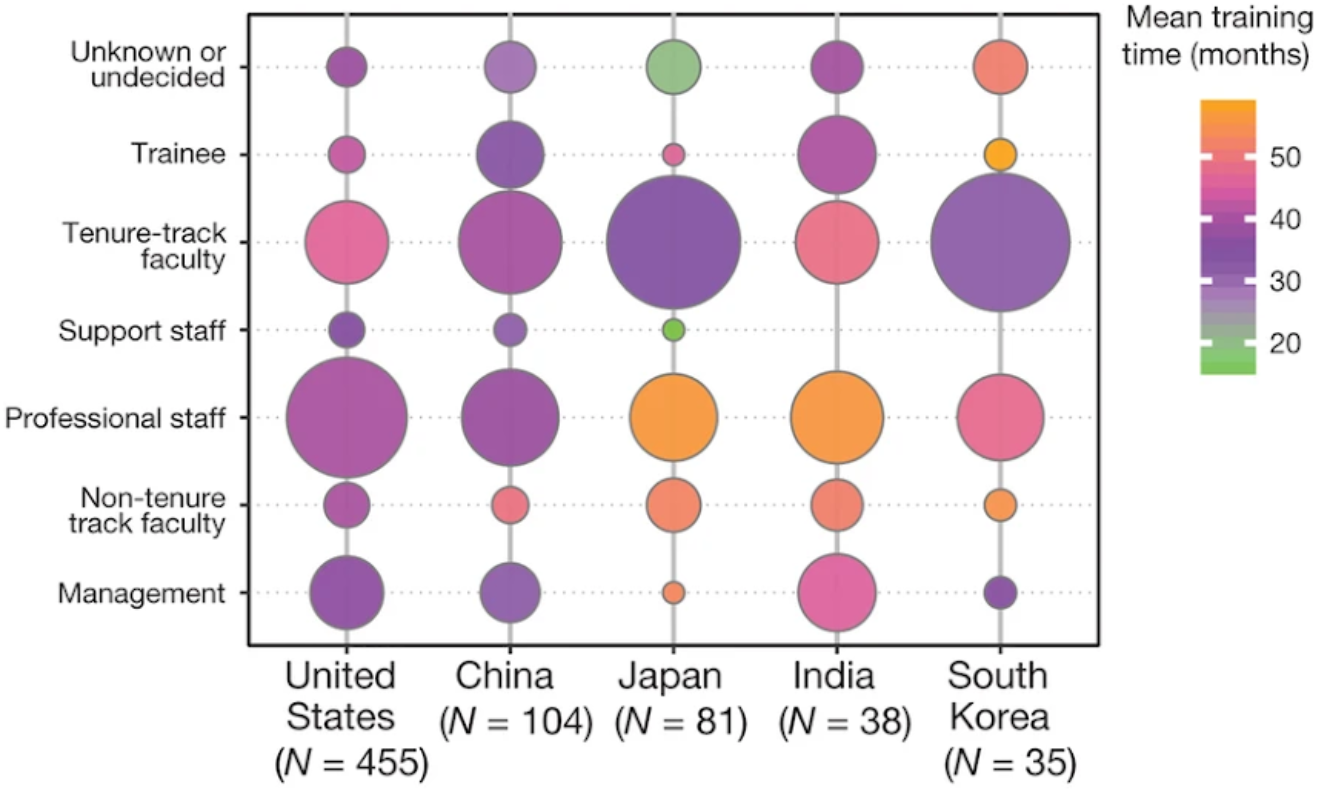
Using a Bubble Plot with an Overlaid Heatmap to Illustrate Career Outcome Differences by Country of Origin, Job Type, and Training Time. A bubble plot showcasing career outcomes from the National Institute of Environmental Health Sciences postdoctoral alumni illustrates how training times vary for those of different countries of origin entering into different job types. The bubble plots illustrate that U.S. alumni enter into professional staff roles at a proportionally higher rate than those from other countries. Alumni from Japan and South Korea are more likely to enter into tenure-track faculty positions than those from other countries. When viewing the heatmap, it becomes apparent that alumni from Japan and India who enter into professional staff positions spend more time in training than those from other countries entering into different job types. (Reproduced without changes; https://creativecommons.org/licenses/by/4.0/)^10^

**Figure 6.**
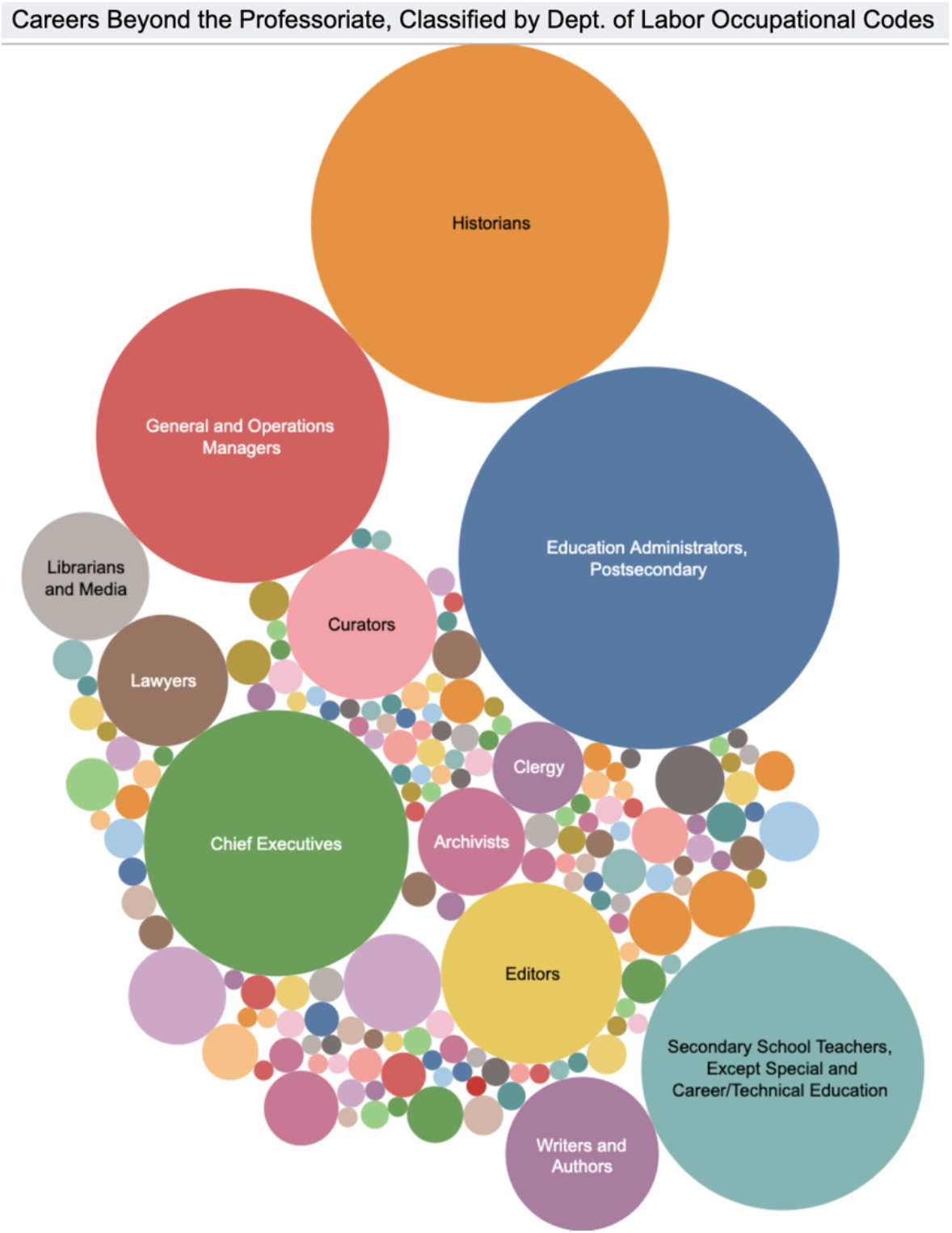
Using a Bubble Plot to Illustrate the Proportion of Individuals Entering into Careers Classified by SOC Codes. The American Historical Association determined the career outcomes of historians and classified them by SOC codes. Their outcomes are depicted in the form of a bubble plot which illustrates the relative proportion entering into different career paths, with the two largest paths being Postsecondary Education Administrators and Historians. Reproduced with permission from AHA.

##### Two-way Table

A two-way table, or contingency table, allows one to visualize the relationship between two sets of categorical variables. In an example from Georgetown University depicting the outcomes of those who earned master’s degrees, one can view the relative proportion of graduates who enter into different sectors, or job functions, as it relates to their job type/industry^27^ (Fig. 7). From this visualization, it is apparent that those entering into academia are primarily conducting research or are in education. Those who enter BioPharma are most likely conducting research, while others may be engaged in regulatory affairs or marketing/communications. Another example of a two-way table can be found in a study from the University of California-San Francisco in an analysis of their postdoctoral alumni outcomes^5^.

**Figure 7.**
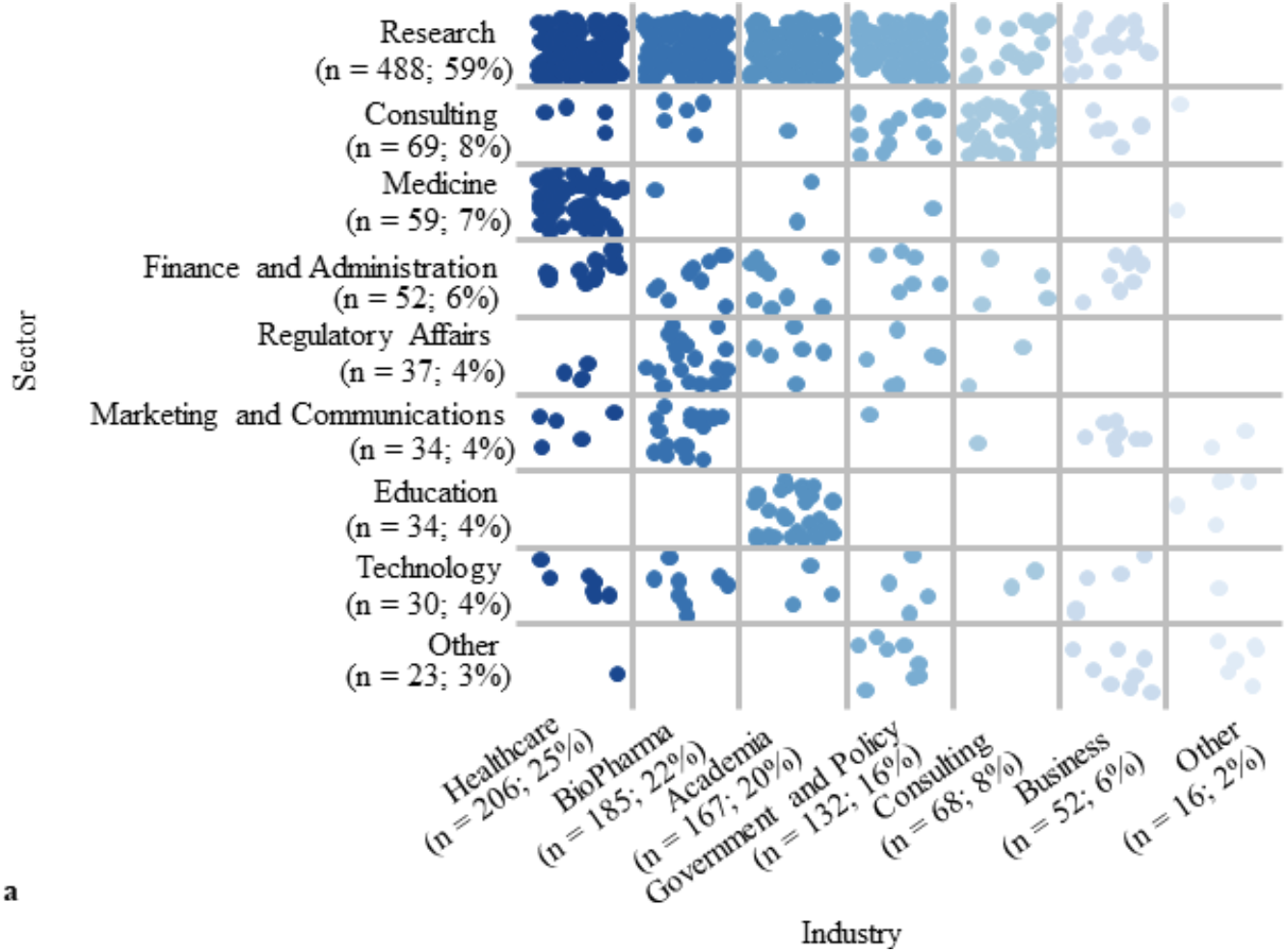
Using a Two-Way Table to Visualize the Career Outcome Relationship Between Sector and Industry. A two-way table was used to illustrate the first destination career outcomes of 829 master’s graduates from Georgetown University within each sector (job function) and industry (job type). An individual graduate is represented as a dot, with the dots color-coded by industry. It is apparent that most graduates are engaged in research in either the healthcare, BioPharma, academia or government and policy industry. (Reproduced without changes; https://creativecommons.org/licenses/by/4.0/)^27^

**Figure 8.**
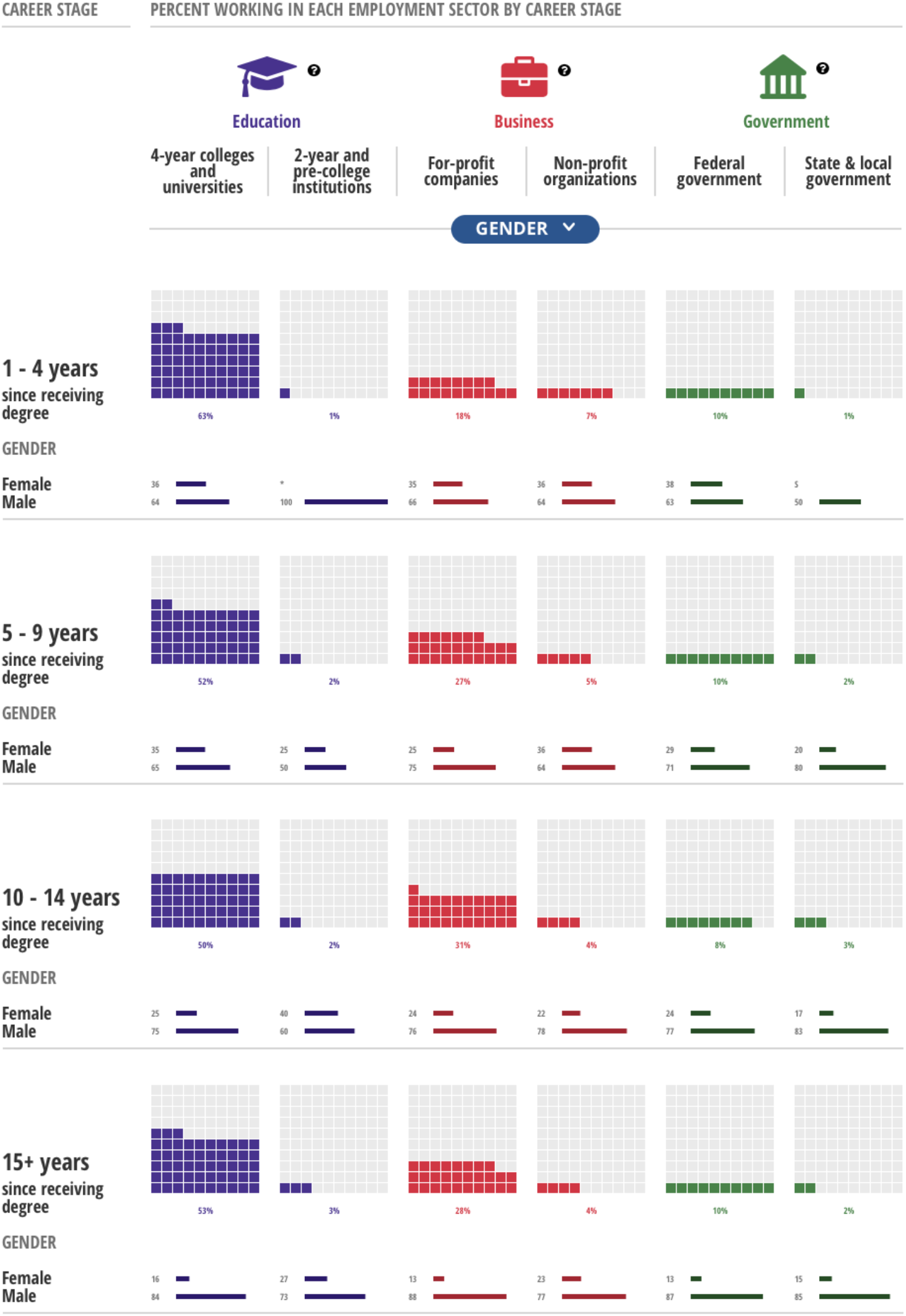
Using Waffle Plots and Small Multiples to Illustrate Career Outcomes and Gender Differences at Different Time Points. The National Science Board used waffle plots used to showcase the relative proportion of science, engineering, and health doctorate (SEH) graduates who entered into different employment sectors relative to how long ago they received their doctoral degree. There is little difference across the various time points, with most entering into 4-year colleges and universities regardless of whether they were 1-4 or 15+ years from receiving their degree. Small multiples represent the relative proportion of males and females comprising the workforce across the different employment sectors and time points. It is apparent that those 15+ years from receiving their degree are largely male. Reproduced with permission from the National Science Board.

**Table 1:** Comparison of Platforms & Visualization Types (adapted from AAMC GREAT 2018 project^28^)

| Platform or Visualization Type | Cost | Time/FTE/Training investment level | Skill Level (Advanced, Intermediate, Novice) | Strengths | Points to Consider | Selected Examples |
| --- | --- | --- | --- | --- | --- | --- |
| **Tableau** | + | + | Intermediate | Easier to create a dashboard by selecting options & less programming literacy required | Less control of data presentation | U Toronto, Stanford, Michigan, etc. |
| **R (& NIEHS dashboard)** | Free | +++ | Advanced | More control of how to present data (predeveloped dashboard can be modified for institutions) | Need programming literacy | NIEHS |
| **Excel** | + | + | Novice | Ease of use for those without much technical experience or resources | Limited visualization options, not much room for customization, not interactive, and low quality | OHSU |
| **Career Services Management Platforms** | +++ | + | Novice | Optimized for those without extensive data analysis experience; systems automate many of the visualizations; some level of customization available | Limited to the visualization options included in the systems with not as much room for customization | Symplicity, Handshake, 12Twenty |
| **Microsoft PowerBI** | + | ++ | Advanced | Easy data sharing and intuitive user-interface; can customize visualizations; visualizations are interactive | Not as widely used in the field and may require additional training; may not contain all visualization options desired | Wayne State, University of Texas System, Weill Cornell Medicine, NACE |
| **Data Analytics Software (Prism, SPSS, SAS)** | Varies (+, +++) | Varies (+, +++) | Advanced | Customizable options for analysis (SPSS, SAS, Prism) and data vis (SAS, Prism) | Plug and play dropdown menus, relatively easy to learn (Prism & SPSS); versatile options by checkbox and extensive tutorials (Prism); coding more time intensive but more customizable (SAS, SPSS) | NIH BEST |
| **Institution-Specific** | +++ | +++ | Advanced | Fully customizable and can make use of available Google or other libraries | Requires programming and software development knowledge; likely requires partnering with groups across one's institution | Boston University, Clemson, UBC |
| **Divergent Stacked Bars** |  | + | Novice | Readily see trends in outcomes that decrease or increase over time at diverging point | More difficult to ascertain changes in bars not precisely at the divergent point; care should be made to explain the X-axis | OHSU, Scripps, NIEHS |
| **Sankey** |  | ++ | Advanced | Show broad patterns of complex data all at once | Sector flow and time flow can be misinterpreted | OHSU, NIEHS, Wayne State, Michigan, UBC, Stanford |
| **Directional Chord/Chord** |  | + | Advanced | Show broad patterns of complex data all at once | Not always immediately obvious; need arrows | NIEHS, Williams College |
| **Stacked Donut/Sunburst** |  | + | Intermediate | Readily see differences in career outcomes across different year-cohorts | Obligate smaller interior donut and larger exterior donut could obfuscate some comparisons | Wayne State |
| **Bubble (+Heatmap)** |  | ++ | Advanced | Compare outcomes of different cohorts simultaneously; overlay with additional dimensions | Relative bubble size; overestimation | NIEHS, American Historical Association |
| **Two-way Table** |  | + | Novice | Compare two sets of categorical variables; illustrate with values, heatmap or dots | Color coding by variables could be mistaken for a heatmap or could obfuscate comparisons | Georgetown University, UCSF |
| **Waffle Plot** |  | + | Novice | Select outcomes can be clearly highlighted as progress towards 100% | Too many outcomes on the same plot could become complicated | National Science Board |
| **Small Multiples** |  | ++ | Intermediate | Powerful method allowing for a quick assessment of overall patterns in data | If multiples are too small they become illegible | National Science Board |

##### Waffle Chart and Small Multiples

A waffle chart allows one to show how the relative proportion of a selected career outcome relates to the whole. The waffle is depicted in the form of a square grid, with the colored areas of the grid representing the data. An excellent example can be found in the National Science Board (NSB) infographic (https://www.nsf.gov/nsb/sei/infographic2/?yr=2013&fd=Mathematics%20and%20statistics&cs=ShowGender) which showcases employment sector data of science, engineering, and health (SEH) doctorates. The data are further represented as small multiples, which is a series of graphs shown together with the same axes and scale allowing for direct comparisons; viewers can thus quickly ascertain how employment trends differ amongst those who received their degrees 15+ years ago versus those who received degrees more recently (relative to when the surveys were conducted in either 1993, 2003, or 2013). The infographic makes further use of small multiples in the form of bar charts to illustrate additional details within each employment sector, including gender, race/ethnicity, job satisfaction, job related to degree, and job duties. For instructions on how to create small multiples in platforms such as Tableau or Excel, multiple websites offering guidance are available online. (https://depictdatastudio.com/data-table-to-small-multiples/ or https://www.juiceanalytics.com/writing/better-know-visualization-small-multiples). Guidance on creating a waffle plot can be found at Depict Data Studio (https://depictdatastudio.com/charts/waffle/) or one can use the waffle plot template from an infographics toolbox from google (https://docs.google.com/drawings/d/1KlcdyB3Pkl3NFH7XyO83yPOBKSXcc7km-4k4shgUlq0/edit?ntd=1).

### 3. RESOURCES, TOOLS, AND COALITIONS

Below, we describe a variety of coalitions and collaborative efforts whose aim is to facilitate others’ ability to collect and disseminate information on the career outcomes of graduate students and postdoctoral scholars. We also describe some key resources (often available for a fee) that can further aid institutions in managing, analyzing, and reporting these data.

#### RESOURCES & TOOLS

The **Institute for Research on Innovation in Science (IRIS)** is a consortium of universities whose members pay to receive access (https://iris.isr.umich.edu/membership/join/) to an IRB-approved data repository that is housed at the University of Michigan. Member institutions share alumni data with IRIS (https://iris.isr.umich.edu/research-data/), which is then de-identified and combined with datasets from other sources (for example, the U.S. Census Bureau, public federal sources, private sector sources, etc.) to obtain a more complete picture of both the career outcomes of alumni as well as the social and economic impacts of investment into the domestic U.S. scientific research enterprise. As a result of domestic focus, many of the repositories used to combine data are only available within the U.S. (with the exception of some Canadian data sources); this therefore precludes the ability to obtain information on international fellows.

**Steppingblocks** is an education and workforce data analytics provider that has entered into a unique partnership with IRIS (https://www.steppingblocks.com). Steppingblocks’ core technology is to structure public data from a multitude of sources, and to employ machine learning to de-duplicate, clean, and categorize data down to the individual level (https://www.youtube.com/watch?v=K6sh0GpOdT4). With the linkage of IRIS and Steppingblocks’ data together, a wealth of information may be obtained, such as award funding history, funding timeframe, employer, salary, etc. This information may then be distributed to members in the form of reports specifically tailored to an institution’s needs. IRIS members (with a membership cost) can also request a download of all underlying data in tabular form so that data obtained from IRIS can be combined with other university-specific datasets. A key aspect of IRIS membership is assistance with publicity to highlight accomplishments of the research enterprise. A press release accompanies each member institutions’ report.

**Academic Analytics** is a company originally created to collect and analyze data on institutions’ research (https://academicanalytics.com). Academic Analytics now also collects graduate and postdoctoral outcomes data. Institutions that hire the company provide them with a list of graduates (names and the year or date of graduation) for Academic Analytics to track, via internet searches, the location of their graduates' employers. Most subscribing institutions provide 10 years of graduate names and contact information and receive collected data for those individuals identified. The company provides person-level information, employer name, position title, and three types of classifications based on the following taxonomies:

- Coalition for Next Generation Life Science (CNGLS) (http://nglscoalition.org/progress/)
- North American Industry Classification Systems (NAICS)/Standard Industrial Classifications (SIC) & Standard Occupational Code (SOC) Listings (https://www.nsca.org/naicssoc-codes/)
- AAUDE (https://www.aaude.org/)

All classifications are done systematically following a logic tree. Data are updated every year, with a rolling 10-year window. Visualizations are simple, and done in Tableau, displaying basic information on career types, race, gender, ethnicity, citizenship, etc.

**Burning Glass Technologies** is a company that, for a fee, provides real-time information on the labor market (such as how it was affected by COVID-19^29,30^) by scanning millions of job ads across the globe and conducting in-depth data analytics to determine what skills are sought after and where certain jobs are located (https://www.burning-glass.com). They recently added “Alumni Analysis”, which is an add-on to their “Labor Insight” platform, to their suite of offerings. With this feature, Burning Glass tracks graduate-level career outcomes by searching their database containing millions of resumes and social media profiles. Based on a snapshot of their alumni outcome platform, Burning Glass allows an institution to select the taxonomy they would like to view, and automatically determines what percentage of alumni are employed within their field of study. The platform also shows the relative geographic location of employment.

**Economic Modeling Specialists International (EMSI)** is similar to Burning Glass in that, for a fee, it also provides real-time information on the labor market. Likewise, it contains an “Alumni Outcomes” service in which an EMSI specialist will collaborate with an institution to track the career outcomes of their alumni by searching over 100 million profiles and resumes aggregated from the web (https://www.economicmodeling.com/). The outcomes are classified by SOC/O*NET code, and EMSI will provide the following: 1) an overview of career outcomes that is filterable; 2) a focused snapshot of employment outcomes in specific programs, 3) a spreadsheet with all available information, and 4) an Excel file with embedded pivot tables. Further assistance with analytics is available, such as benchmarking to national standards or allowing students to explore potential wage earnings as a function of career outcome.

Several **Tools for Automating or Crosswalking Standard Job Codes** have been developed which allow matching of large-scale text, especially from job postings data, to SOC codes. This includes the NIH’s Standardized Occupation Coding for Computer-assisted Epidemiological Research or SOCcer tool (https://soccer.nci.nih.gov/soccer/)^31^. This tool has also been used to analyze sub-sectors of the job market by cross-referencing data scraped from job posting aggregation sites like Indeed.com^32^.

Other tools exist which can “crosswalk” between CIP codes and SOC codes (https://nces.ed.gov/ipeds/cipcode/Files/IES2020_CIP_SOC_Crosswalk_508C.pdf); and between the US Department of Labor’s SOC codes and international codes such as ISCO-08 (https://www.bls.gov/soc/isco_soc_crosswalk_process.pdf), allowing international comparisons. A similar international comparison tool has been developed in Canada, Codage Assisté des Professions et Secteurs d’activité (https://ssl3.isped.u-bordeaux2.fr/CAPS-CA/Langue.aspx). A tool from the CDC can automate assignment of NAICS or SOC codes as well as crosswalk between different versions of each (https://csams.cdc.gov/nioccs/Default.aspx).

#### COALITIONS

**NIH Broadening Experiences in Scientific Training (NIH BEST) Consortium, Rescuing Biomedical Research (RBR), and Future of Bioscience Graduate and Postdoctoral Training (FOBGAPT)** spearheaded efforts to develop, refine, and adopt a unified, 3-tiered career outcomes taxonomy divided by workforce sector, career type, and job function, later named the Unified Career Outcomes Taxonomy and broadly implemented by the member institutions of CNGLS^33^ (http://rescuingbiomedicalresearch.org/rbr-actions/improving-transparency-ph-d-career-outcomes/).

The **Coalition for Next Generation Life Science (CNGLS)** formed in 2017 to address calls for greater transparency in graduate and postdoctoral training. Since formation, coalition membership has expanded from 10 founding institutions to 55 as of June 2021. Member institutions agree to publish career outcomes data for PhD and postdoctoral alumni according to the top two tiers of the UCOT 2017 taxonomy—workforce sector and career type (http://nglscoalition.org).

The **Council of Graduate Schools (CGS)** launched a PhD Career Pathways project to help institutions understand the professional aspirations and career pathways of PhDs. As of February 2021, 75 Doctoral Institutions have participated in the project. Several “Research in Brief” articles have been published, highlighting aggregate results from the survey. (http://cgsnet.org/understanding-career-pathways)

The **American Association of Universities (AAU)** began an initiative in 2014 to encourage institutions to collect student employment outcomes data. They have since formally launched the AAU PhD Education Initiative, whose goal is to make data about PhD career pathways and employment trends widely available. The AAU has also collated and summarized lists of career outcome tracking efforts (https://www.aau.edu/sites/default/files/AAU-Files/PhD/10.18.18_Multi-Institutional_Efforts.pdf and https://www.aau.edu/sites/default/files/AAU-Files/PhD/10.18.18_Multi-Institutional_Efforts.pdf and https://www.aau.edu/sites/default/files/AAU-Files/PhD/Project-Summaries-02.22.19-1.pdf).

**The National Academies of Science, Engineering, and Medicine (NASEM)** released two consensus reports in 2018 about graduate and postdoctoral STEM education and training: “The Next Generation of Biomedical and Behavioral Sciences Researchers: Breaking Through” (https://www.nap.edu/catalog/25008/the-next-generation-of-biomedical-and-behavioral-sciences-researchers-breaking) and “Graduate STEM Education for the 21st Century”. (https://www.nap.edu/catalog/25038/graduate-stem-education-for-the-21st-century) The first study made a key recommendation for institutions to collect and disseminate career outcomes data using common standards and definitions. It suggests the NIH incentivize such data collection by making it a requirement. The second study recommended transparency of career outcomes in order for current and prospective students to be able to make educated choices and to enable institutions to make effective adjustments. It also urged the need for standardization, transparency, and accessibility.

**Future of Research** members contributed to the NASEM report on the biomedical research enterprise (Breaking Through), and participated in a workshop on mentoring (The Science of Effective Mentorship in STEM) with discussions incorporated into the NASEM report. The group has also organized workshops and built a webpage, “Tracking Career Outcomes at Institutions” (http://futureofresearch.org/tracking-career-outcomes-at-institutions/), collating a list of U.S. institutions and organizations that have collected and published career outcomes data.

The **American Association of Medical Colleges (AAMC)** released the National M.D.-PhD Program Outcomes Study in April 2018 (https://store.aamc.org/downloadable/download/sample/sample_id/162/), covering the career paths of physician-scientists. AAMC representatives also worked with RBR and NIH BEST to adopt a unified career outcomes taxonomy. AAMC **Group on Graduate Research Education and Training (GREAT Group)** additionally organizes annual conferences to promote and discuss graduate education, professional development, and training related topics including career outcomes tracking.

**National Institute of General Medical Science (NIGMS)** issued an updated Institutional Predoctoral Training Grant Funding Opportunity Announcement in 2017, which features new guidelines for predoctoral training grant (T32) applications. Among other changes, applicants will need to provide information about career outcomes, career exploration opportunities, and professional skills development for their trainees (https://loop.nigms.nih.gov/2017/10/new-nigms-institutional-predoctoral-training-grant-funding-opportunity-announcement/ and https://grants.nih.gov/grants/guide/pa-files/PAR-17-341.htmlx).

The **National Institute of Environmental Health Sciences (NIEHS)** extramural division developed a system entitled ‘CareerTrac’ (https://careertrac.niehs.nih.gov/public/staticPage/about) to enable tracking of trainees’ employment outcomes and accomplishments over time from laboratories receiving NIH funding. It was the first system of its kind to be developed at an NIH institute, and is also used by the Fogarty International Center, NIGMS, National Cancer Institute, and the National Institute of Diabetes and Digestive and Kidney Diseases (NIDDK). A poster describing outcomes from those previously on T32 training grants has been presented (https://www.niehs.nih.gov/research/supported/assets/docs/a_c/careertrac_evaluating_t32_training_outcomes_508.pdf).

**NORC at the University of Chicago: Progress and Pitfalls in Monitoring Doctoral Degree Holders’ Career Paths** is an NSF grant-funded project that supports four main activities: a web-based national survey of graduate deans in fall 2018 to assess current practices of monitoring graduates’ careers; a set of focus groups of graduate deans in December 2018 that will address guiding questions informed by the survey; a one-and-a-half-day conference in May 2019 with the goal of developing standards for collecting and reporting data on doctoral career pathways; and a multipronged dissemination of the project results (https://reports.norc.org/white_paper/identifying-promising-practices-in-university-based-monitoring-of-doctoral-career-pathways/).

The **American Psychological Association (APA)** has partnered with Economic Modeling Specialists International (EMSI) for tracking and analyzing the post-graduation outcomes of psychologists, which they expect will provide a wealth of information useful for creating career development resources, as well as determining the impacts of the coronavirus pandemic on the psychology workforce. In this study, APA also aims to identify the specific skill sets that psychologists are using in their careers. At present, the APA’s Center for Workforce Studies has a wealth of information on the psychology workforce, such as the one found on their data tools outlining the types of activities that psychologists are engaged in (https://www.apa.org/workforce/data-tools/careers-psychology).

The **Graduate Career Consortium (GCC)’s Outcomes Committee** collated information on the publicly available graduate and postdoctoral alumni career outcomes from each of its member institutions. As of September 2020, approximately 63% of member institutions publicly report on their career outcomes in a quantitative manner. A comprehensive database of these outcomes was released (https://osf.io/28dn6/) and will be updated annually each fall^12^.

#### CONCLUSION

Tracking, compiling, and publishing PhD career outcomes are crucial to the future of the biomedical research enterprise and higher education, and are part of growing appeals for systemic change (NIH, NIH BEST, Council of Graduate Schools, Graduate Career Consortium, National Academies, Coalition for Next Generation Life Science, Rescuing Biomedical Research, etc.). Efforts to gather and share career outcomes data can help inform training and education initiatives related to the decline of available faculty jobs^34^, economic and workforce needs, evolving postdoctoral policies, institutional accountability, and growing diversity in the career paths available to PhD recipients^35^. The maintenance of career outcomes data is thus becoming an essential practice for institutions that issue PhDs, and standardization of such practices will: 1) empower prospective students and postdoctoral scholars to make informed decisions; 2) enable effective intra-institutional planning and decision making; 3) facilitate inter-institutional comparisons and benchmarking; and 4) enable institutions to evaluate their ability to address equity, diversity, and inclusion (e.g. how do gender or race and ethnicity affect outcomes, and how does an institution’s training environment and structure equitably support all students/postdocs as they navigate into careers?^26,36,37^).

Our review aims to support systemic change by providing an overview of key classification systems complete with highlights and caveats of these systems, which is accompanied by the creation of a crosswalk tool that maps similarities between each system. In addition, we provide a synopsis of methods for visualizing PhD career outcomes, as well as tools for automating or outsourcing the tracking of this data, thus enabling institutions to choose from a variety of system(s) that work best for them. Transforming graduate and postdoctoral training and shaping the future of the research enterprise and broader economy is reliant upon informing our educators, the workforce, and our trainees about the many impactful career options pursued by graduate students and postdoctoral scholars.

## ACKNOWLEDGEMENTS

This project was a collaborative effort by members of the Graduate Career Consortium (GCC) PhD Outcomes Committee. The authors would like to thank for their critical review and helpful comments on the manuscript: Rebecca Dunning and Laura Stark; Gina Shereda, Michael Tessel, and Brandy Simula. We thank Reinhart Reithmeier, Rosanne Lurie, Anne Laughlin/Susi Varvayanis, and Jackie Wirz for their contributions towards the University of Toronto 10,000 PhDs, NACE taxonomies, Cornell’s methodology of tracking and visualizing outcomes, and general visualizations, respectively. Thanks to Allison Fryer, Amanda Mather, and Jackie Wirz (Oregon Health & Science University) and Nadine Lymn (National Science Board) for permission to showcase career outcomes. Thanks to Lisa Barge for sharing a compiled list of classification systems from public and private institutions.

